# Biocontrol of mushroom crop mycoparasites by novel *Bacillus velezensis* strains

**DOI:** 10.1101/2025.09.02.673639

**Authors:** William Kay, Jaime Carrasco, Souvik Kusari, Marjon Krijger, M. José Carpio, Thomas Barnes, M. Sonia Rodríguez Cruz, Jan van der Wolf, Till Bebenroth, Gail M. Preston

**Affiliations:** University of Oxford, Department of Biology, South Parks Road, Oxford, OX1 3RB, United Kingdom; IRIAF – Agri-food and Forestry Regional Research and Development Centre, CIAF - Agroforestry Research Centre, Carretera Toledo-Cuenca km 174, Cuenca 16194, Spain; Centro Tecnológico de Investigación del Champiñón de La Rioja (CTICH), Autol, Spain; Institute of Natural Resources and Agrobiology of Salamanca, Cordel de Merinas 40-52, 37008 Salamanca, Spain; Department of Genetics, Institute of Biology and Ecology, Faculty of Science, Pavol Jozef Šafárik University in Košice, Mánesova 23, 04154 Košice, Slovakia; Center for Mass Spectrometry, Faculty of Chemistry and Chemical Biology, Technische Universität Dortmund, 44227 Dortmund, Germany; Wageningen University and Research, Biointeractions & Plant Health, 6700 AA Wageningen, the Netherlands; Institute of Agricultural Sciences, ICA-CSIC, Serrano 115 bis, 28006 Madrid, Spain

**Keywords:** *A. bisporus*, Mycoparasites, Casing material, Biocontrol, Secondary metabolites, Specialised metabolites, Lipopeptides, *Bacillus velezensis*, lipopeptides, fengycin, *Agaricus bisporus*

## Abstract

1.

The cultivation of button mushroom (*Agaricus bisporus*) requires the design of tailor-made substrates that nourish the crop and promote morphology changes from mycelium to basidiome. The agronomic stages of mushroom development are also influenced by the microbiota present in the mushroom crop microcosm, which may have a beneficial impact on mushroom growth, development and quality, or a detrimental impact through reduction of yield or quality as parasites, competitors or disease vectors in mushroom crops. In this report we describe the isolation of multiple strains of *Bacillus velezensis* from mushroom casing material and basidiomes that show antifungal activity towards major mushroom mycoparasites, along with further characterization of their mode of action. Full genomes of *B. velezensis* CM5, CM19, CM35, EM5 and EM39 were sequenced and annotated, which together with metabolic profiling of specialised metabolites produced by CM5, CM19 and CM35 suggested that the antifungal activity of these strains is linked to the production of the lipopeptide fengycin. However, in crop trials, these strains did not increase mushroom yield or provide significant control of the mushroom parasite *Lecanicillium fungicola*. Genomic and analytical tools were designed and used to evaluate *B. velezensis* persistence in casing when the selected strains were artificially applied. *B. velezensis* population levels decreased significantly after application, potentially contributing to the lack of biocontrol activity observed in crop trials.

**Key Points:**

1. *B. velezensis* strains isolated from peat-based microcosms are shown to have significant inhibitory effects against major fungal parasites of mushrooms *in vitro*.
2. Through full genome sequencing and metabolic profiling using LC-HRMS, the antifungal activity is correlated with the production of lipopeptides, particularly fengycin analogues.
3. In trials using button mushroom crops artificially infected with *Lecanicillium fungicola*, the causal agent of dry bubble, treatment of casing material with these strains did not significantly limit dry bubble disease or increase yield.
4. The persistence of strains in crop when artificially applied showed significant decreases after application on casing, potentially contributing to the lack of biocontrol activity.

## 4. Introduction

The commercial cultivation of the white button mushroom (*Agaricus bisporus*) relies on a 3-phase composting process followed by a bio-based casing layer that provides the cues required for fructification (Dias et al. 2021). Fungal parasites such as green mould (*Trichoderma* spp.), wet bubble (*Mycogone perniciosa*), dry bubble *(Lecanicillium fungicola*) and cobweb (*Cladobotryum* spp.) cause major crop losses (Gea et al. 2021). Breeding for resistance is difficult (Sonnenberg et al. 2017), and key fungicides including chlorothalonil, carbendazim, metrafenone and prochloraz-Mn are now prohibited or ineffective (Commision 2019; Kosanović et al. 2013), necessitating alternative control measures. Alongside hygiene, biocontrol is a promising component of integrated disease management (Gea et al. 2021), operating through mechanisms such as antagonism via enzymes, antimicrobial metabolites or VOCs, competition, growth stimulation and enhanced nutrient uptake (Carrasco and Preston 2020; Zhang and Sun 2018).

*Bacillus velezensis* includes well-studied strains with strong biocontrol activity mediated by antimicrobial metabolites (Kenfaoui et al. 2024; Rabbee et al. 2023), including bacillaene, bacillibactin, bacilysin, difficidin, fengycin, macrolactin H and subtilin (Pandin et al. 2018b). Fengycin, of particular relevance for suppressing fungal parasites, is synthesised by a non-ribosomal peptide synthetase (NRPS) complex encoded by the *fen* operon (*pps* in B. subtilis) (Desmyttere et al. 2019; Gao et al. 2018; Zihalirwa Kulimushi et al. 2017), and disrupts fungal membranes via sterol interactions (Mantil et al. 2019). Fengycin-producing strains inhibit *Trichoderma aggressivum* (Kosanovic et al. 2021; Pandin et al. 2019), *Cladobotryum mycophilum* (Clarke et al. 2022a; Clarke et al. 2024), *L. fungicola* (Clarke et al. 2022b) and *M. perniciosa* (Novikova and Titova 2023).

Fengycin variants include classes A and B (Steller et al. 1999) and additional analogues such as fengycin S and C (Sang-Cheol et al. 2010; Villegas-Escobar et al. 2013), whose production in *B. amyloliquefaciens* is positively regulated by *comQXPA*-mediated quorum sensing (Yin et al. 2024).

In this study, we isolated several novel *B. velezensis* strains showing broad antifungal activity from casing materials and basidiomes. We characterised these strains via phylogenetics and comparative genomics, evaluated three isolates for in-crop antimicrobial activity through artificial inoculation, assessed their persistence on casing and impact on the native microbiota, and identified secreted specialised metabolites using LC-HRMS to determine correlations with antifungal activity.

## 5. Methods

### 5.1 Isolation of the cultivable bacterial microbiome

Cultivable bacteria were isolated from three commercially used mushroom casing materials: black peat (peat moss-based; Euroveen B.V., BVB Substrates, Grubbenvorst, The Netherlands), blonde peat (sphagnum peat moss-based; Valimex KF, Valencia, Spain), and a 50:50 mixture of both materials.

The physical and chemical characteristics of these casings have been reported previously (Carrasco et al. 2019). Cultivable bacteria were also isolated from *A. bisporus* basidiomes following the method of Aslani *et al*. (2018).

#### Microbiome cultivation

i) for casing-colonising bacteria, 10 g of casing was suspended in 100 mL of 0.1% sterile bacteriological peptone broth (Thermo Scientific #LP0037B); ii) for basidiome bacteria, healthy closed mushroom caps were submerged in 1% NaClO for 30 s and rinsed with sterile distilled water. 10 g of internal mushroom cap tissue was homogenized (pestle and mortar) and suspended in 100 mL of 0.1% bacteriological peptone broth. Samples were incubated in an orbital shaker-incubator (Ovan 1000001087, Badalona, Spain) at 120 rpm and 25 ºC for 10 min. The supernatant was serially diluted and spread on peptone agar. Media recipes in Table S1.

#### Isolation and preservation of individual colonies

One hundred and forty-one individual colonies (Table S2) were re-isolated to plates containing Luria Bertani (LB) Agar and incubated at 28 °C for 24-48 h. Single colonies were added to LB and incubated between 25 and 28 °C (150 rpm) between 5 and 18 hours. Fermentation was stopped when the absorbance at OD_600_ reached a value of 1. The resulting cultures were re-suspended in 20% glycerol solution and stored at -80ºC (Panasonic MDF-U5386S-PE) until further use.

### 5.2 DNA extraction, sequencing, transformation

#### DNA extraction and Sanger sequencing of 16S rDNA

Single colonies were grown in LB (28 °C, 220 rpm, 24 h), pelleted, washed, and DNA extracted using the NucleoSpin Plant II kit (Macherey-Nagel). The 16S rDNA region was amplified using primers 27F/1492R (Stackebrandt and Goodfellow 1991) and Phusion Master Mix (Thermo Fisher, UK) with standard cycling conditions. PCR products were treated with ExoSAP-IT™ and Sanger sequenced by Eurofins (Luxembourg) or Source Bioscience (Nottingham, UK). Sequences were aligned using MAFFT and taxonomically identified via BLASTn searches against the NCBI database (Camacho et al. 2009).

#### Full genome sequencing and analysis

Isolates were resuspended in 0.5 mL of DNA Shield (Zymo Research) and sequenced by MicrobesNG. Raw reads were trimmed with Trimmomatic 0.30 with a quality cutoff of Q15 (Bolger et al. 2014), assembled with SPAdes version 3.7 (Bankevich et al. 2012), and annotated using Prokka 1.11 (Seemann 2014). Genome visualisation used Artemis (Rutherford et al. 2000), assembly metrics were assessed with Quast (Gurevich et al. 2013), and taxonomic assignment was performed using Kraken (Wood and Salzberg 2014). Additional annotations were generated via RAST/SEED (Aziz et al. 2008; Aziz et al. 2012) and comparative genomics conducted with NCBI PGAAP (Tatusova et al. 2016). Biosynthetic gene clusters were predicted with antiSMASH (Blin et al. 2021). All genome sequences are deposited in NCBI under BioProject PRJNA1142391. Details of quality Assessment of *B. velezensis* genomes using QUAST can be found in Table S3.

#### Phylogenetic analysis of *B. velezensis* strains

Five housekeeping genes (*gyrB, pgk, rpoB, rpoD, tuf*) were amplified using the NucleoSpin Plant II kit (MACHEREY-NAGEL GmbH & Co. KG) and primers listed in Table S4, then sequenced (Source Bioscience). Homologous sequences from comparative strains were retrieved from NCBI. Concatenated alignments were generated using MUSCLE (Edgar 2004), and phylogenies inferred with IQ-TREE using 1000 bootstrap replicates (Nguyen et al. 2015). Trees were visualised with iTOL (Letunic and Bork 2021).

#### Transformation of *B. velezensis* CM19 with pHAPII gfp+

The plasmid pHAPII gfp+ (Cao et al. 2011) was extracted from a transformed strain of *B. velezensis* QST713 (kindly provided by Romain Briandet, INRAE) using the NucleoSpin Plasmid extraction kit (Macherey-Nagel). Subsequently, *B. velezensis* CM19 cells were transformed using a room temperature electroporation protocol (Morales-Ruiz et al. 2019).

### 5.3 *In vitro* fungal inhibition assays

#### *In vitro* confrontation trials

Four fungal parasites: *T. aggressivum* (TAV1), *M. perniciosa* (M25), *C. mycophilum* (CM13900) from La Rioja mushroom farms, and *L. fungicola* (150/1) from Warwick HRI (Banks et al. 2019), were cultured on potato dextrose agar (PDA) for 3–7 days and resuspended in sterile water. Bacterial strains were grown overnight in LB. For dual-culture assays, 10 µL droplets of fungal and bacterial suspensions were placed 30 mm apart on PDA and incubated at 25 °C in darkness for 4–13 days depending on the parasite. Biocontrol activity was quantified by measuring inhibition zones.

#### Spore germination assays

Fungal spores were harvested from PDA using sterile water, filtered through Miracloth, and counted by haemocytometer. Aliquots (100 µL; 107–108 spores mL^-1^) were spread onto PDA plates and dried before spotting 10 µL of an overnight culture of *B. velezensis* CM19 pHAPII-gfp at the centre. Plates were incubated at 28 °C for 1-4 days, after which agar sections spanning the bacterial colony and inhibition zone were excised, mounted on slides, and stained with propidium iodide (PI; 1 µL mL^-1^). Samples were imaged by confocal microscopy (Leica TCS SP5) using GFP (Ex 488 nm; Em 510–530 nm) and PI channels (Ex 561 nm; Em 625–645 nm). Identically treated, unchallenged fungi served as controls.

#### Effect of isolated bacteria on *A. bisporus* mycelium growth

*A. bisporus* H15 was cultured on 2% agar compost medium (prepared from powdered compost extract) for 7 days in darkness. Bacterial overnight cultures (10 µL) were spotted 3 cm from the fungal inoculation point, with four droplets per plate (one per quadrant). After a further 7 days of incubation, photographs were taken and the radial distance between bacterial droplets and the fungal colony edge was measured. LB-only droplets served as controls.

#### Toxicity of bacteria to *A. bisporus* sporocarp tissue

Ten mm cubes of mushroom tissue were inoculated with bacteria from overnight LB cultures at a rate of 100 µL per cube (LB only as control). Cubes were photographed after 48h and assessed for browning using a 0-4 scale where: 0 was no effect; 1 = light scarring; 2 = light browning; 3 = heavy browning; and 4 = black/dark brown with obvious seepage.

### 5.4 Biocontrol activity of *B. velezensis* in a crop trial infected with *L. fungicola*

#### Bacterial inoculum preparation

Bacterial strains were grown overnight in LB, and cultures with OD600 > 0.7 were used to inoculate a 5 L Biostat A fermenter (Sartorius, Germany). Liquid fermentation was carried out at 28 °C, pH 7, and 10-50% dissolved oxygen for 7-8 h until OD600 ≈ 1. Cells were harvested by centrifugation (4500 rpm, 10 min, 10 °C), washed twice with PBS (phosphate buffer, pH = 7.4), and resuspended in 10% skim milk-10% sucrose as cryoprotectant. The suspension was frozen, freeze-dried, pulverised into soluble powder, and viable counts were confirmed by serial dilution plating on LB.

#### Crop Trial

An experimental growth chamber (IGCS 1500 HR LED, Ibercex) was used following commercial mushroom-growing practices. The trial used 32 randomized blocks (0.04 m2 each), each containing 1250 g Phase III compost (Sylvan H15 strain supplemented with Mylo Pro and Champfood) and 750 g black casing (CNC, Valimex, or Euroveen). Six treatments (Table S5) included water controls ± *L. fungicola*, three *B. velezensis* strains (CM5, CM19, CM35) + *L. fungicola*, and a fungicide control (Prochloraz-Mn). Prochloraz-Mn (1 g m^-2^) was applied on day 4 post-casing; bacterial inocula on day 5; and *L. fungicola* 150/1 on day 10. *L. fungicola* conidia (10^6^ conidia m^-2^) were applied in 20 mL of 2 × 10^6^ conidia L^-1^ (Gea et al. 2014). Bacterial treatments consisted of 20 mL of 2 × 109 cfu L^-1^ (final 10^9^ cfu m^-2^). Disease incidence (healthy and diseased mushrooms and bubbles) and biological efficiency of the crop were assessed at the end of the first and second flushes.

#### Analysis of PLFAs

Casing was sampled at four key stages: raw casing (day 0), fully colonised casing (day 10), fructification of the first flush (day 21), and end of the first flush (day 29). Samples were analysed for phospholipid fatty acids (PLFAs) following (Frostegård et al.) (1993). Total PLFAs were used as a proxy for microbial biomass (nmol g^-1^), and specific biomarkers quantified: Gram-negative (monounsaturated fatty acids and cyclopropyl 17:0), Gram-positive bacteria (iso and anteiso saturated branched chain fatty acids), Actinobacteria (10-methyl fatty acids) and fungi (18:2 ω6 cis and 16:1w5) (Zelles 1999). Fatty acid methyl esters (FAMEs) were analysed by gas chromatography.

#### Statistical Analysis

PLFA-derived total bacterial and fungal biomass was analysed via ANOVA after Levene’s tests for homogeneity. Post-hoc comparisons used Tukey or Games–Howell depending on variance structure (p < 0.05). Analyses were conducted in SPSS v24 (SPSS Inc. Chicago, USA), and principal component analysis (PCA) was performed using PAST v3.15 (Hammer et al. 2001).

### 5.5 TaqMan assays for *B. velezensis* strains (CM19, CM5 and CM35)

#### DNA extraction

Bacteria were collected with a loop from the agar surface of 90 mm diameter tryptic soy agar (TSA) plates into a 2 mL tube and used for DNA extraction. The Wizard Genomic DNA purification kit (Promega) was used for DNA extraction. DNA yield was determined using the Qubit ds DNA HS assay kit with the Qubit Fluorometer (Life Technologies).

#### TaqMan design

Potential target sites were identified using the whole chromosomal genome sequence and CLC genomic workbench (Qiagen, Aarhus, DK). Target sequences were dissected in 500 bp-long sequences and mapped to *B. velezensis* QST713 (CFSAN0334339, CLC mapping settings: length fraction: 0.85, similarity fraction: 0.85, global alignment: no). Candidate regions were checked for sequence similarity with non-target organisms using BLASTn (NCBI) (Camacho et al. 2009). At least six sets of primer/probe combinations were designed per target on 500 bp target-specific fragments using primer quest tool of Integrated DNA Technologies (IDT, Leuven, Belgium) with default settings. Two sets of primer/probe combinations were selected for the design of the final triplex TaqMan assay, targeting different genes, based on the kinetics. The specificity of the assays was tested using genomic DNA of 27 strains (Table S6). Full methods in Methods S1.

#### Triplex TaqMan design

The two simplex assays for the target bacteria were combined with an assay that quantifies *Xanthomonas campestris* pv. *campestris* (Xcc) into a triplex TaqMan assay (Table S7, Methods S2). Methods for the limit of detection are found in Methods S3.

### 5.6 Surfactant activity and lipopeptide production

#### Surfactant production

*B. velezensis* strains were cultured overnight in Optimised Medium (OM, Table S1) and standardised to OD_600_ = 1. Droplets (10 µL) were pipetted onto the lid of a Greiner CELLSTAR® 96 well plate, allowed to sit for 2 min, and measured using an eye piece graticule on a ZEISS dissecting microscope. Mean droplet diameters of 3 experiments, each containing 2 droplets, were assessed for differences against OM only control.

#### Lipopeptide fraction collection

Lipopeptides were isolated from 600 mL CM19 cultures incubated in OM at 30 °C for 24 h. After centrifugation (10,000 g, 10 min), supernatants were acidified to pH 2.0 with concentrated HCl and precipitated overnight at 4 °C. Pellets were collected by further centrifugation (4 °C, 15 min), resuspended in methanol, and shaken for 2 h. Extracts were filtered (0.4 µm), concentrated to 1 mL by evaporation, and applied to filter discs for confrontation assays.

#### Extraction of agar plates for targeted metabolic profiling

Five LB agar plates per strain (CM5, CM19, CM35) were grown for 5 days at 28 °C. Biomass was pooled, homogenised, and extracted sequentially with 50 mL of HPLC-grade MeOH using an ultrasonic bath. Extracts (after 10 min) were filtered, evaporated to dryness at 35 °C (using rotary evaporator), and re-dissolved in 1 mL MeOH. After centrifugation (8000 rpm, 5 min), samples were analysed by HPLC-HRMS.

#### High-performance liquid chromatography-high resolution mass spectrometry (HPLC-HRMS)

Extracts were analysed using an LTQ Orbitrap XL mass spectrometer (Thermo Scientific, USA) equipped with a HESI-II source and coupled to an Agilent 1200 HPLC system (pump, PDA detector, column oven, autosampler). Separations were performed on a Luna C18(2) column (50 × 3 mm, 3 µm; Phenomenex), with an Aeris Peptide XB-C18 column (150 × 2.1 mm, 3.6 µm) used for HRMSn analyses. Mobile phases consisted of H_2_O (+0.1% formic acid; A) and CH_3_OH (+0.1% formic acid; B) at a flow rate of 350 µL min^-1^. The gradient programme was: 5% B for 2 min; 5–100% B over 24 min; 100% B for 6 min; return to 95% A in 0.5 min and re-equilibration for 3 min. For targeted fengycin HRMSn measurements, the gradient was modified to 5% B for 2 min; 5–90% B over 8 min; 90–100% B over 20 min, followed by re-equilibration as above. The mass spectrometer was operated in positive ion mode at a resolving power of 60,000 (m/z 400), scan rate 1 Hz, and mass range m/z 160–1600. N-butyl benzenesulfonamide ([M+H] ^+^, m/z 214.08963) was used as lock mass. Nitrogen served as sheath and auxiliary gas, helium as collision gas, and HRMSn spectra were acquired using collision-induced dissociation at 42 eV. An authentic fengycin standard (≥90%; Sigma-Aldrich) was used for confirmation. Data were processed using Xcalibur v2.2 SP1.48, with masses filtered by intensity (I > 1.0 × 10^3^) and a maximum mass tolerance of 2 ppm. Background subtraction was applied where required.

## 6. Results

### 6.1 Novel *B. velezensis* display antimicrobial activity

From 141 isolated strains (Table S2), *Bacillus* spp. and *Pseudomonas* spp. were the most abundant taxa (Figure 1A). Initial confrontation assays (Table S8) identified *B. velezensis* strains as the most antagonistic against the four major fungal parasites of *A. bisporus* (*T. aggressivum, M. perniciosa, L. fungicola*, and *C. mycophilum*) and these were selected for further analysis (Figure 1B). Nine *B. velezensis* strains, along with *Bacillus oceanisediminis* CM26 as a negative control, were evaluated *in vitro*; all exhibited significant inhibition relative to the control (Figure 2A, 4B; ANOVA: *C. mycophilum* p < 0.001; *L. fungicola* p < 0.001; *M. perniciosa* p = 0.139; *T. aggressivum* p < 0.001; post-hoc Tukey, p < 0.05). The interaction of all four parasites with CM19 pHAPII gfp+ was visualised using propidium iodide (PI) staining to monitor membrane integrity and cell viability. All four pathogens were frequently non-viable or produced germ tubes/hyphae that became PI-positive or lysed (Figure 2C–E). Most *B. velezensis* strains showed minimal antagonism toward *A. bisporus* itself and caused little to no browning of sporocarp tissue (Figure 2F), supporting their suitability as biocontrol candidates.

**Figure 1.**
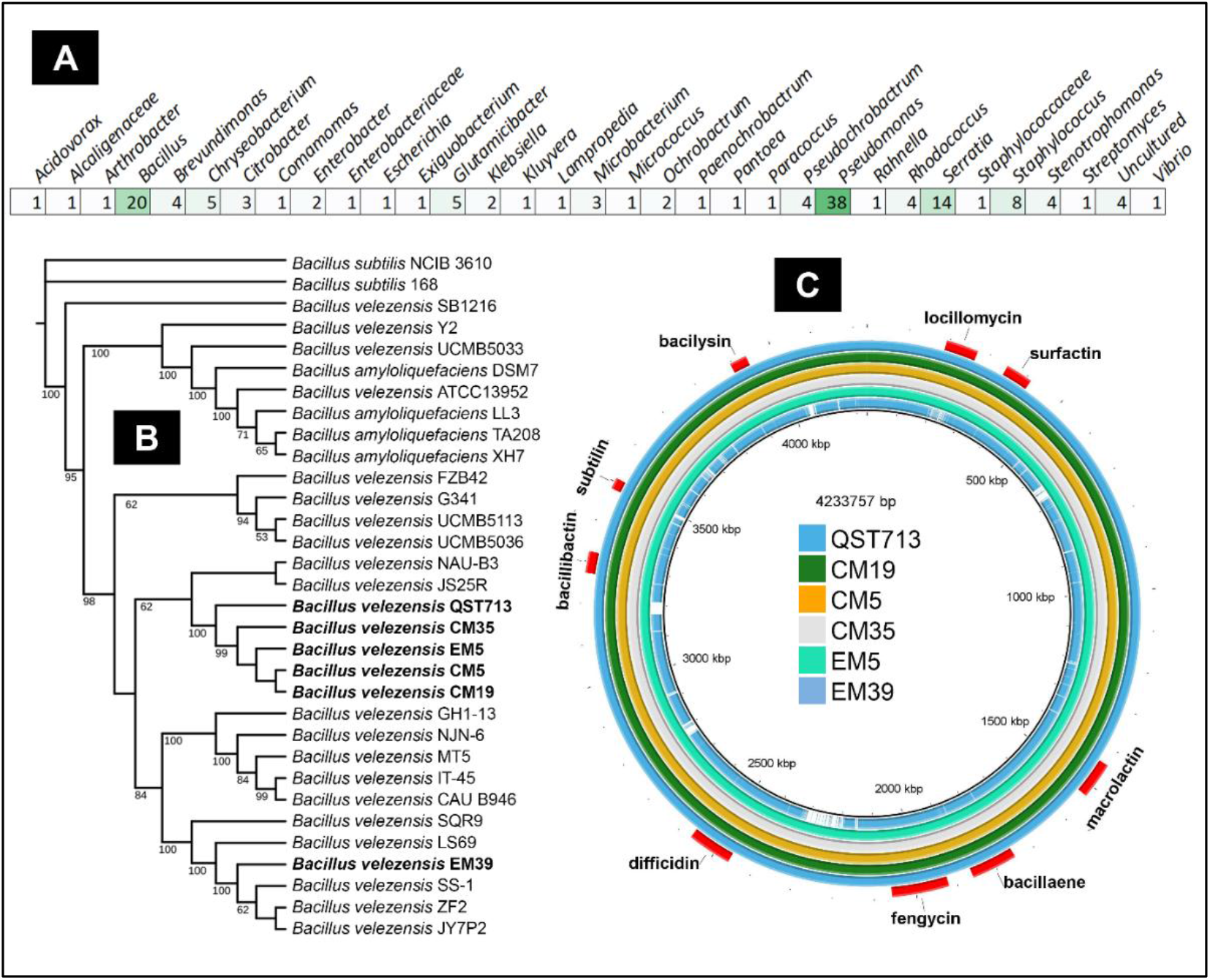
A) Cultivable bacterial isolates from three types of casing layer and endofungal bacteria from basidiomes of *A. bisporus* identified using 16S rDNA sequencing; B) Phylogenetic tree of selected *Bacillus* strains built using concatenated genomes of five housekeeping genes (*gyrB, pgk, rpoB, rpoD, tuf*) aligned to reference strain *B. velezensis* FZB42. Strains highlighted in bold were novel to this study; C) Visualization of *B*. genomes using BLAST Ring Generator (BRIG) (Alikhan et al. 2011); Putative metabolite biosynthetic clusters compared to QST713 are shown by red marks in outer ring.

**Figure 2.**
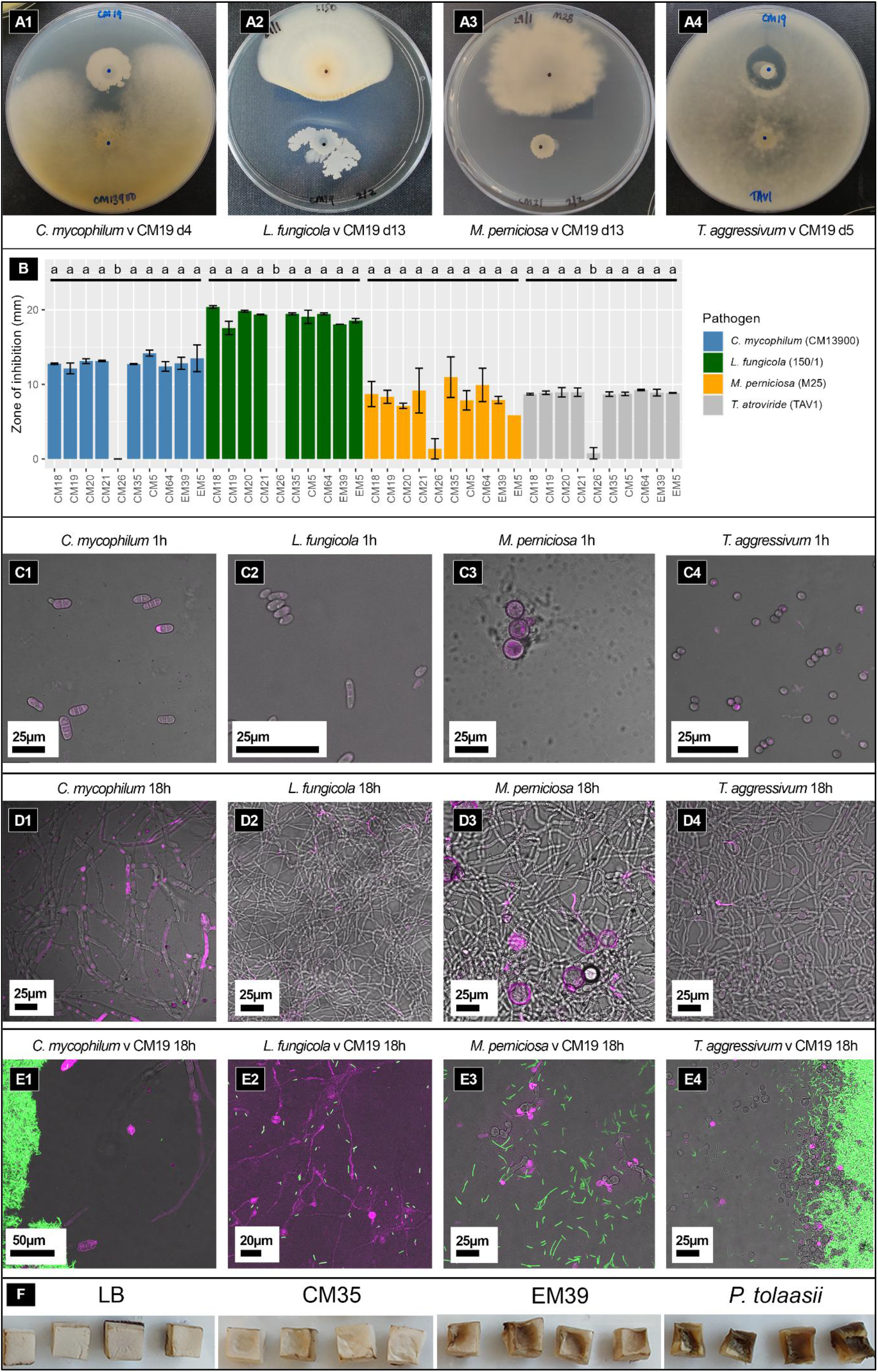
Antifungal activity observed in *B. velezensis* strains: A) Antagonism between *B. velezensis* CM19 and the four fungal parasites (d = days after inoculation). B) Graph shows the mean (+/-SE) fungal inhibition zone (mm) of the three biological replicates. Letters indicate statistical differences (one-way ANOVA and Tukey’s HSD test α = 0.05). C & D) Unchallenged fungal parasites grown *in vitro* (PDA plates) observed using confocal microscopy: t=0 (C), and t=18 h (D). E) Confocal images of fungal parasites confronted with *B. velezensis* CM19 pHAPII gfp (green) *in vitro*. Live dead staining with propidium iodide (PI; pink) (C, D, E). F) Sporocarp cubes 24h after inoculation with bacterial strains, *P. tolaasii* NCPPB2192 (positive control) or LB (negative control).

### 6.2 Mushroom casing isolates of *B. velezensis* are closely related to the commercial biocontrol strain *B. velezensis* QST713

Phylogenetic analysis of five *B. velezensis* isolates together with the commercial strain QST713 resolved three distinct clades, *B. amyloliquefaciens, B. velezensis*, and *B. subtilis*, consistent with previous classifications (Fan et al. 2017; Pandin et al. 2018a) (Figure 1B). Four isolates (CM5, CM19, CM35, EM5) clustered tightly within the *B. velezensis* clade and showed close relatedness to QST713. Genome sequencing of five casing isolates (CM5, CM19, CM35, EM5, EM39) confirmed this relationship, with four strains sharing 99.99% nucleotide identity with QST713 based on NCBI PGAAP analysis (Table S9). A complete list of SNP-containing genes and predicted amino-acid changes in these strains relative to QST713 is provided in Table S10.

### 6.3 Fengycins are the most abundant anti-microbial lipopeptides produced by casing isolates of *B. velezensis in vitro*

antiSMASH analysis identified multiple biosynthetic gene clusters (Table 1) with homology to those encoding surfactin, fengycin, bacillibactin, bacilysin, and the antimicrobial polyketides bacillaene, difficidin and macrolactin. Surfactin and fengycin are known lipopeptides with antimicrobial and surfactant properties (Gilliard et al. 2024; Sur 2019), whereas bacillibactin functions as an iron-chelating siderophore (Qin et al. 2019).

**Table 1:** Comparative antiSMASH analysis of gene clusters predicted to encode genes involved in specialised metabolite synthesis between *B. velezensis* CM5, CM19, CM35, EM5, EM39 and QST713.

| Most similar cluster | Type | CM5 | CM19 | CM35 | EM5 | EM39 | QST713 |
| --- | --- | --- | --- | --- | --- | --- | --- |
| bacillaene | Polyketide+<sup>1</sup>NRP | 100% | 100% | 100% | 100% | 100% | 100% |
| bacillibactin | NRP | 100% | 100% | 100% | 100% | 100% | 100% |
| bacilysin | Other | 100% | 100% | 100% | 100% | 100% | 100% |
| butirosin A/B | Saccharide | 7% | 7% | 7% | 7% | 7% | 7% |
| difficidin | Polyketide | 100% | 100% | 100% | 100% | 100% | 100% |
| fengycin | NRP | 100% | 100% | 100% | 100% | 100% | 100% |
| locillomycin | NRP+Polyketide | 28% | 28% | 28% | 28% | - | 28% |
| macrolactin H | Polyketide | 100% | 100% | 100% | 100% | 100% | 100% |
| subtilin | <sup>2</sup>RiPP: Lanthipeptide | 100% | 100% | 100% | 100% | - | 100% |
| surfactin | NRP:Lipopeptide | 82% | 82% | 82% | 82% | 82% | 82% |

<sup>1</sup>NRP: Non-Ribosomal Peptide (built by non-ribosomal peptide synthetases (NRPSs)); <sup>2</sup>RiPP: Ribosomally synthesized and post-translationally modified Peptide.

Quantification of surfactant production by droplet diameter assays (Akpa et al. 2001) showed that all *B. velezensis* strains exhibit significantly increased droplet sizes compared with the LB-only control (Figure S1), consistent with secretion of surface-active metabolites.

Targeted HPLC-HRMS profiling of CM5, CM19 and CM35 detected fengycin (1×CO, 1×OH, Ala:R1=13C) in all strains under the tested conditions (Figure 3). The distinction between fengycin A and B substitution of alanine (A) or valine (B) at position six was confirmed via comparison with an authentic standard and literature-based fragmentation patterns (Ma et al. 2016; Wang et al. 2004).

**Figure 3.**
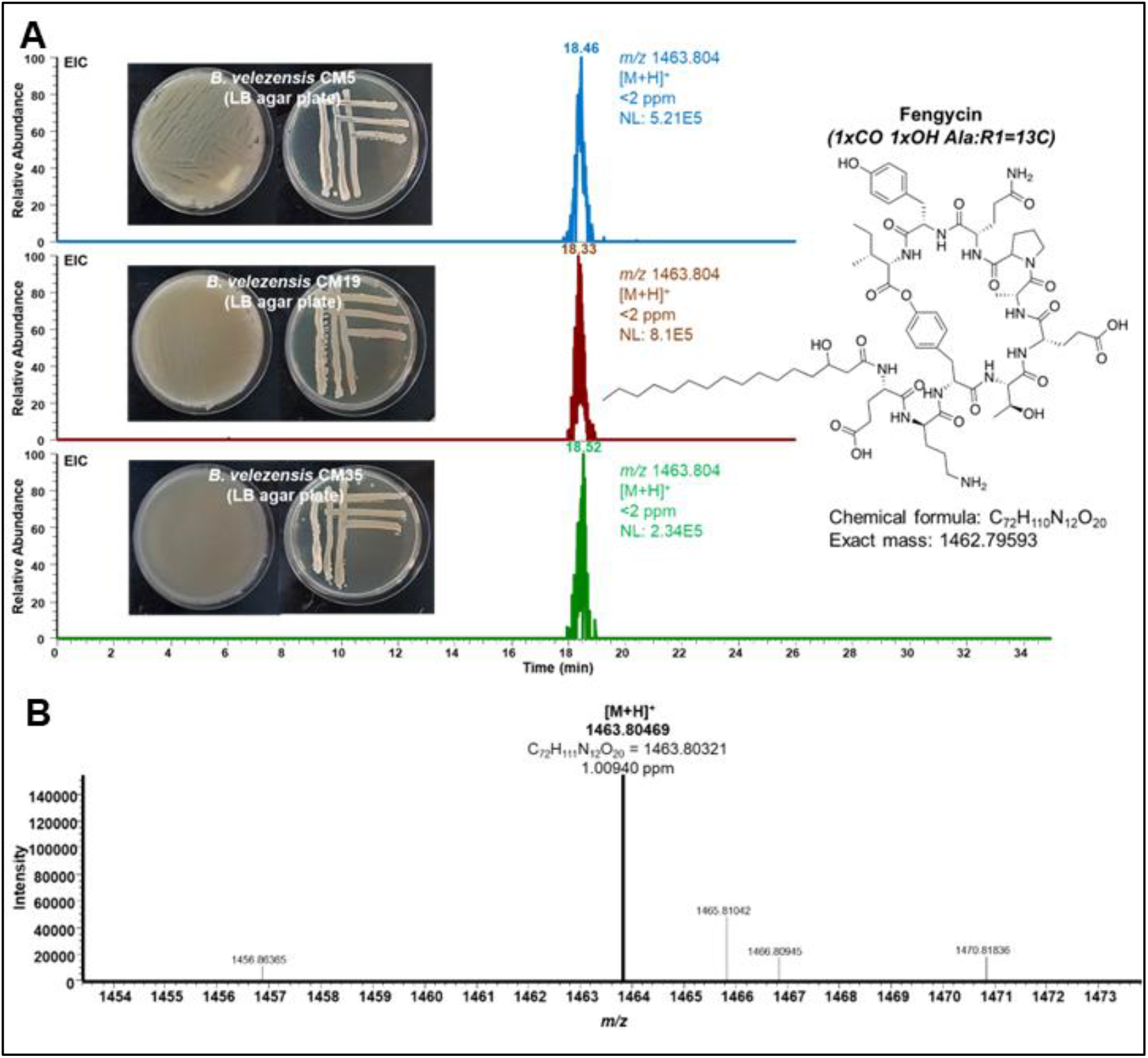
HPLC-HRMS analysis of fengycin produced by *B. velezensis* strains CM5, CM19 and CM35 cultivated on LB agar. (A) Extracted ion chromatograms ([M+H]^+^, *m/z* 1463.804, ±2 ppm) show the production of fengycin (highest physiological amounts: CM19> CM5> CM35). Phenotypic characteristics of the strains grown on LB agar (left) and the chemical structure of fengycin is shown (right). (B) HRMS spectrum of fengycin produced by the bacteria.

Other predicted metabolites, including macrolactins (Table S11), were not detected under the tested conditions (i.e., < limit of detection (LOD)).

### 6.4 The application of *B. velezensis* strains to mushroom crops was ineffective in limiting disease caused by *L. fungicola*

Three *B. velezensis* strain (CM5, CM19, CM35) were taken forward for crop trials designed to assess (i) effects of the treatment on the biological efficiency of the crop, and (ii) biocontrol effects against the fungal parasite *L. fungicola*. Disease was successfully established following inoculation with LF and statistically significant differences between control groups (+LF and –LF) were detected (Figure 4A), confirming that artificial inoculation with LF affected yield. The control inoculated with LF (Figure 4A) was the least productive in terms of biological efficiency, and treatment with a single dose of CM19 was the most productive. However, differences between control and *B. velezensis* treatments were not found to be significant (*p* > 0.05) in the first flush, second flush or total cropping cycle (Figure 4A). As expected, the use of prochloraz-Mn was found to significantly reduce disease in both the first and second flush.

**Figure 4.**
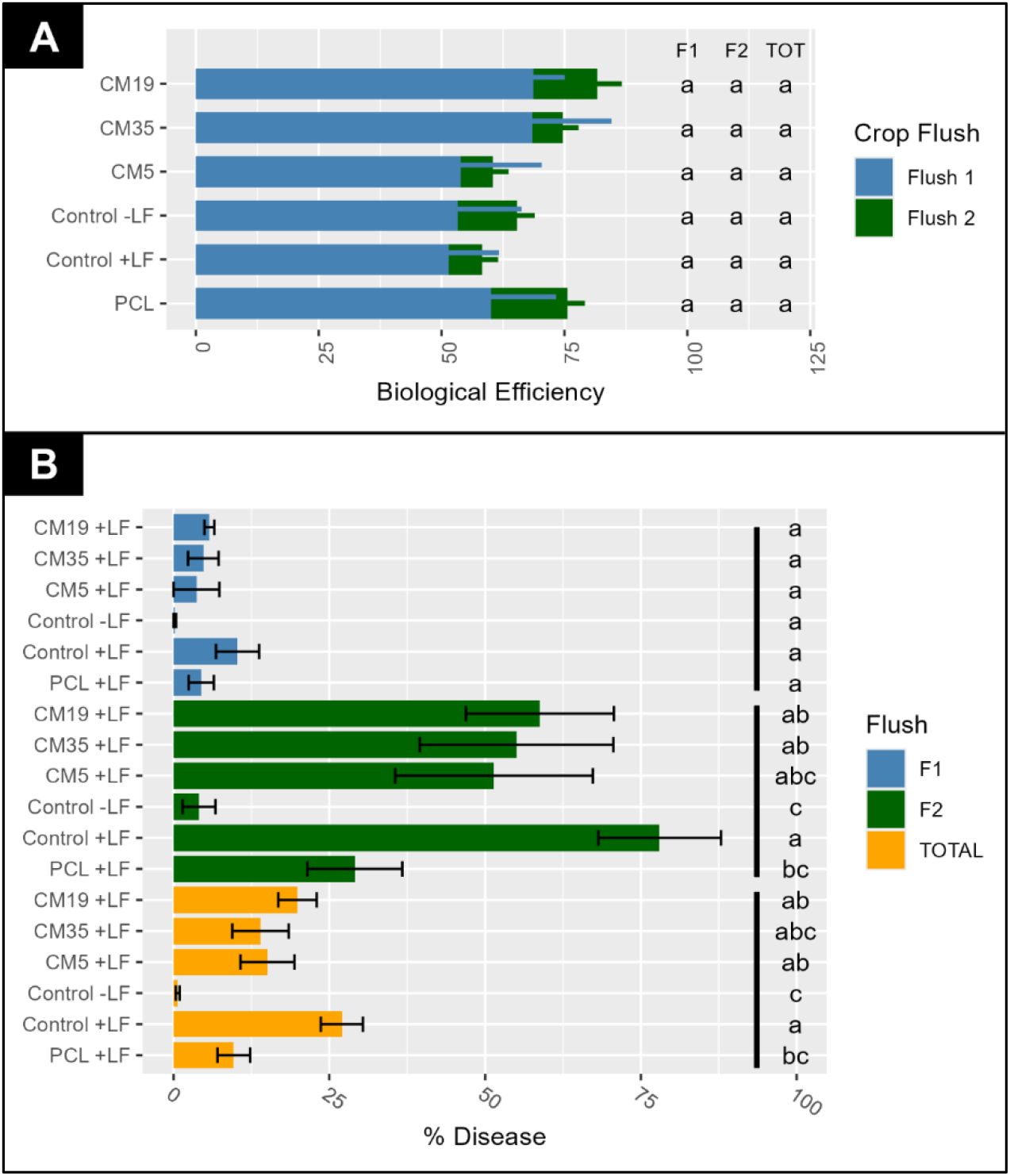
Crop trial of *B. velezensis* vs *L. fungicola* 150/1. A) Biological efficiency (BE) of the crop (Kg of mushroom per 100 Kg of compost in dry weight) at flush 1 (F1), flush 2 (F2) and for the total cropping cycle (TOT); B) Diseased mushrooms (%) collected from F1, F2 and for the total cropping cycle. Graphs show the mean (+/-SE) of 3 biological replicates. Letters indicate groups of statistically different conditions (one-way ANOVA and Tukey’s HSD test α = 0.05). +LF / -LF = inoculated with / without *L. fungicola*. PCL = Prochloraz-Mn.

### 6.5 Phospholipid fatty acid analysis (PLFA)

The impact of *B. velezensis* supplementation on casing microbial communities was evaluated using PLFA profiling to quantify microbial biomass and assess community structure across key developmental stages of the crop trial. PLFA profiling across crop stages showed a significant increase in total microbial biomass over time. In control treatments, total PLFA increased 3.4, 5.8, and 9.5-fold at mycelial penetration, first-flush primordia formation, and first-flush mushroom growth, respectively, relative to raw casing (Figure 5A). The relative contribution of bacterial PLFAs declined as cropping progressed, while fungal PLFAs increased and became dominant by day 29.

**Figure 5.**
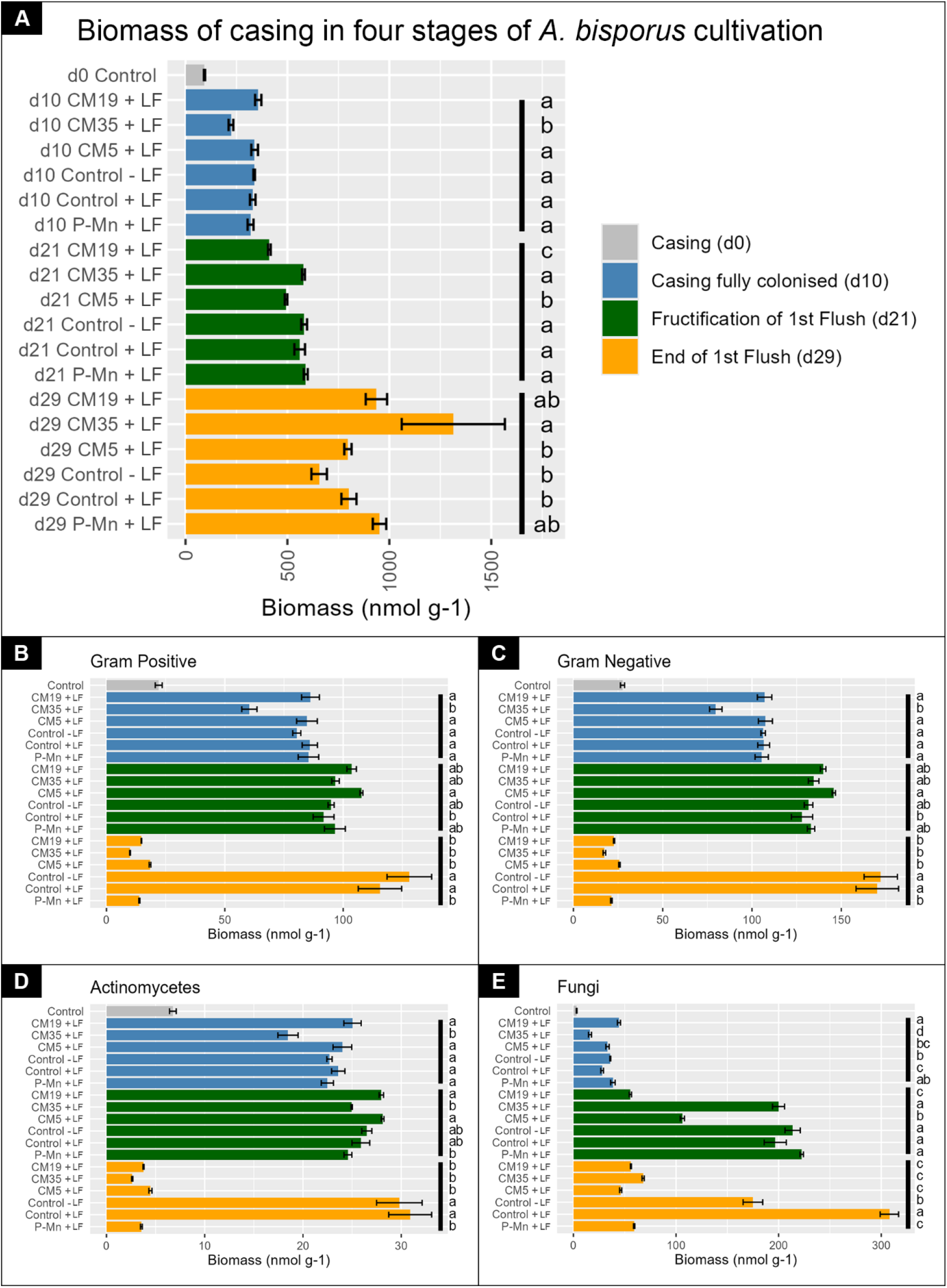
A: Quantification of casing microbial biomass by analysis of PLFAs. Four timepoints were assessed: day 0 (d0) (Beginning of trial; grey), d10 (Fully colonized casing, blue), d21 (Fructification of first flush, green), and d29 (End of first flush, orange). Treatment 1: Control +LF (+); Treatment 2: Control - LF (-); Treatment 3: CM5 +LF; Treatment 4: CM19 +LF; Treatment 5: CM35 +LF; Treatment 6: Prochloraz-Mn (P-Mn +LF). Letters highlight differences between treatments at the same time point (Tukey Test, p > 0.05). Error bars show SE. B: Abundance of Gram-positive bacteria. C: Abundance of Gram-negative bacteria. D: Abundance of Actinomycetes. E: Abundance of fungal PLFAs. All data shown uses nmol g^-1^. LF = *L. fungicola*.

Gram-negative bacterial PLFAs were consistently more abundant than Gram-positive PLFAs across all stages, with both groups most prevalent early in the cycle (Figure 5B–C). Actinomycete-associated PLFAs remained significantly lower than other bacterial groups throughout (Figure 5D).

### 6.6 TaqMan assays for *B. velezensis* indicate low persistence of biostimulants following supplementation to mushroom crops

To determine whether the limited biocontrol effect observed in crop was linked to the persistence of *B. velezensis* in the casing, a subgroup-specific triplex TaqMan assay was developed to quantify supplemented populations. The two *B. velezensis* subgroup assays detected the three target strains as well as CM18, CM20, CM21 and EM5, all members of the same clade (Table S6, Table S7), but did not amplify other *B. velezensis* isolates such as EM39. When applied to casing samples, supplemented *B. velezensis* was detectable only on the day of application (days 1 and 18 after casing; Figure 6A, B). Sensitivity tests showed detection limits of 10^4^cfu g^-1^ in exponential-phase cultures and 10^7^cfu g^-1^ in stationary-phase cultures (Figure 6C), suggesting that limited recovery from casing may reflect low DNA extraction efficiency rather than complete loss of bacterial populations.

**Figure 6.**
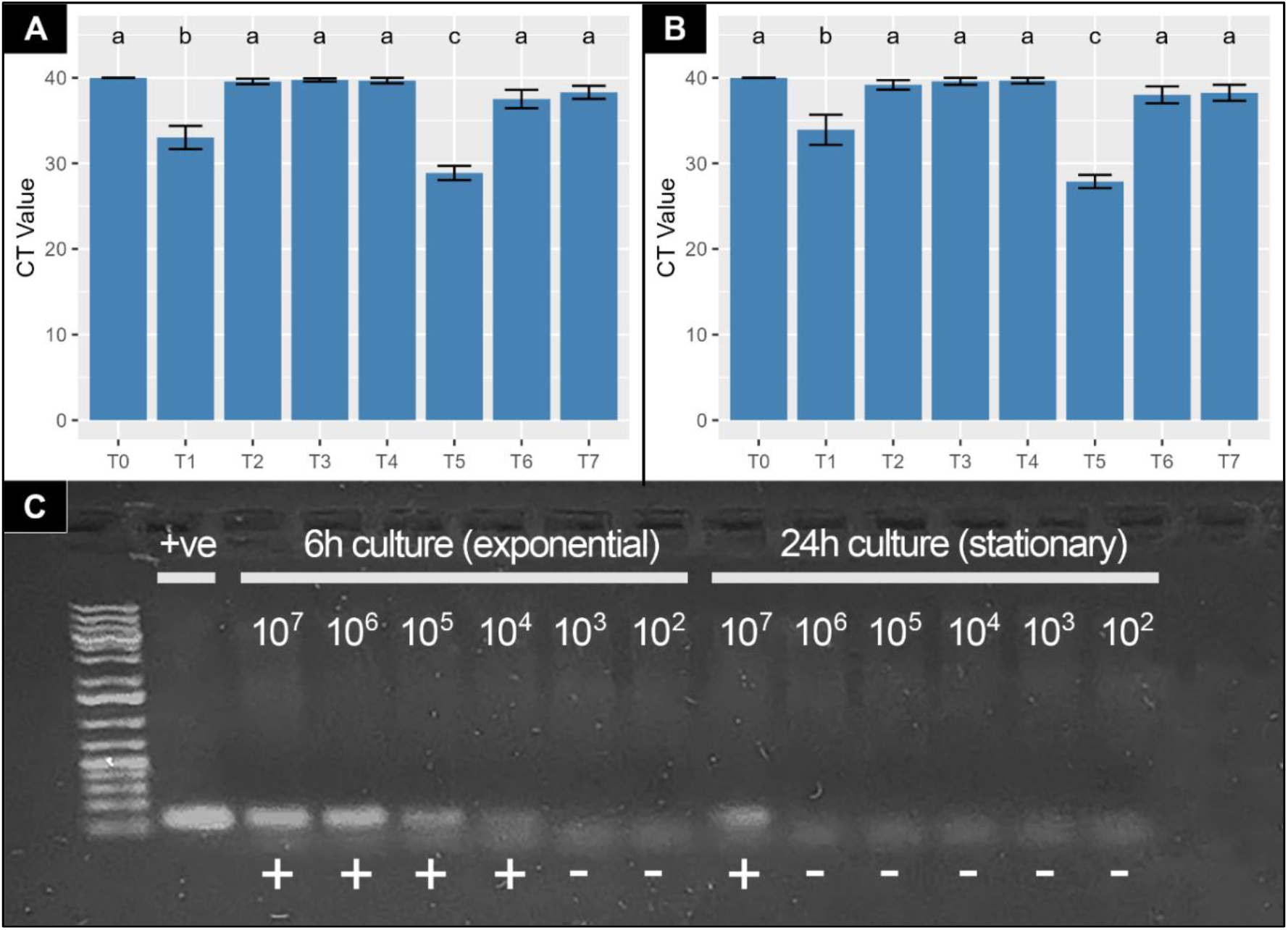
Population dynamics of *B. velezensis* CM19 following application to mushroom casing as determined by a quantitative TaqMan assay. T0 (day 0 after casing), T1 = after application of *B. velezensis* CM19 (day 3), T2 = End of spawn run (day 9), T3 = Beginning of the 1^st^ flush (Day 14), T4 = End of the 1st flush (Day 17), T5 = Directly after the second application of *B. velezensis* CM19 (Day 18), T6 = Beginning of the 2^nd^ flush (Day 22), T7 = End of the 2^nd^ flush (Day 25). A: Results using the HP2.21 (FAM) TaqMan probe; B) Results using the HP2.19 (Cy5) TaqMan probe. C) Electrophoresis gel showing results from PCR reactions using HP2.19 TaqMan primers (35 cycles) for DNA extracted from casing soil supplemented with *B. velezensis* CM19 cells (from LB medium) in either exponential growth stage (6h) and stationary cultures (24 h). Numbers indicate the number of bacteria added to the casing soil before extraction (cells added g-1 of soil).

## 7. Discussion

Button mushroom cultivation involves several agronomic stages (compost preparation, spawning, casing with non-sterile peat-based material and fruiting), each influenced by complex microbial interactions (Carrasco et al. 2021). The casing layer, a microbiologically rich environment essential for fructification and regulated in part by beneficial bacteria (Braat et al. 2022), is also the primary infection site for destructive mycoparasites (Carrasco et al. 2021; Gea et al. 2021). Consequently, the structure and activity of the casing microbiota are central to both mushroom development and disease outcomes. This study aimed to isolate and characterise naturally occurring bacteria from casing soil and basidiomes, assess their antifungal activity against economically important mycoparasites, explore the mechanisms underlying this antagonism, and evaluate their potential as biocontrol agents under crop conditions.

Several *B. velezensis* strains isolated from commercial casing (CM5, CM19, CM35) and from mushroom basidiomes (EM5, EM39), including those described in a published patent (Carrasco and Preston 2023), showed strong antifungal activity *in vitro* against four major mushroom pathogens (*C. mycophilum, L. fungicola, M. perniciosa, T. aggressivum*). This antagonism aligns with previous reports of *B. velezensis* strains functioning as effective biocontrol agents, such as strain Kos from Irish casing soil (Clarke et al. 2022a; Clarke et al. 2022b; Kosanovic et al. 2021) and the commercial product QST713 isolated from peach orchard soil in California (Anastassiadou et al. 2021). Notably, CM5, CM19 and CM35 displayed low toxicity toward *A. bisporus* and did not inhibit mycelial growth at the application rate tested (10^9^ cfu m^-2^). However, despite their *in vitro* antagonism, none of the strains provided effective control of dry bubble disease caused by *L. fungicola* under crop conditions compared with the chemical fungicide control (PCL) (Figure 4B).

Previous studies have similarly reported limited in-crop efficacy of commercial *B. velezensis* QST713 (Serenade®) and *B. amyloliquefaciens* D747 (Amylo-X^®^) against wet bubble, particularly under high inoculum pressure (Navarro et al. 2023). Likewise, QST713 failed to control cobweb disease in artificially infected trials, whereas culture filtrates of *B. velezensis* Kos achieved only moderate efficacy (30–40%) (Clarke et al. 2024). These findings suggest that pathogen species, inoculum density and overall disease pressure strongly influence biocontrol outcomes. In our study, a relatively high inoculum load of *L. fungicola* was used, which may have exceeded the suppressive capacity of the supplemented strains. Improved control might be achievable using lower pathogen pressure, higher or optimised doses of *B. velezensis*, or formulations that enhance antimicrobial compound production before and after application. However, preliminary assays indicated that very high inoculum levels (>10^10^ cfu m^-2^) can negatively impact *A. bisporus* colonisation (unpublished), highlighting the need to refine dose, timing and formulation to balance efficacy with crop safety.

Selective TaqMan assays were developed to track *B. velezensis* dynamics in crop, with two assays per target combined with an extraction/amplification control enabling specific detection in multiplex set-up. However, the limit of detection (LOD) in casing soil (≤4 × 10^5^ cells mL^-1^) restricted accurate monitoring to situations where supplemented populations remain relatively high. Detection efficiency was also reduced for stationary-phase cultures, likely reflecting poor DNA extraction from spores.

When the TaqMan assays were used for samples from the crop trial, supplemented strains were detectable only on the day of application, suggesting either rapid population decline or a transition to sporulation that renders the assay less sensitive. Since *B. velezensis* must adapt to a complex environment, optimizing the formulation or antimicrobial activity of *B. velezensis* based products is key to improve stability, shelf-life and deliver biocontrol activity (Kenfaoui et al. 2024).

Consistent with previous studies (Carrasco et al. 2020; Carrasco et al. 2019), we found that the casing microbiome undergoes marked structural changes during mushroom cultivation. PLFA profiling showed that microbial community composition shifted significantly across developmental stages, with Gram-negative bacteria dominating early colonisation, supporting their recognised role in primordia initiation. Total microbial biomass was highest at the end of fructification, reflecting the strong influence of *A. bisporus* mycelial growth on casing microbiota. Increases in both bacterial and fungal biomass correlated with the progression of host mycelium during the crop cycle, in agreement with earlier work (Carrasco et al. 2020; Wei-Ming et al. 2009). The expansion of bacterial populations may be driven by metabolites released during fruiting body development, which modify nutrient availability and shape microbial community diversity in the casing (Zhao et al. 2022).

Application of *B. velezensis* strain CM5 resulted in increased Gram-negative and Gram-positive PLFA signatures in *L. fungicola-*infected samples at day 21 (fructification of the first flush), suggesting that beyond direct antagonism of mycoparasites, *B. velezensis* may enhance mushroom development indirectly by promoting beneficial bacteria. Such effects could arise through suppression of competing microbes or nutrient release following lysis of susceptible organisms. The persistent dominance of Gram-negative bacteria across all cropping stages supports earlier reports that these taxa, particularly *Pseudomonas* spp., are key stimulators of sporophore initiation, likely through metabolism of inhibitory volatiles in the casing layer (Noble et al. 2003; Rainey 1989; Wei-Ming et al. 2009). In contrast, actinomycetes remained consistently low, likely reflecting their sensitivity to environmental fluctuations (temperature, pH, moisture) during cultivation (Akond et al. 2016; Lacey 1997) and competitive exclusion by faster-growing groups such as *Pseudomonas* spp. (Noble et al. 2003; Rainey 1989).

All five *B. velezensis* strains (CM5, CM19, CM35, EM5, EM39) exhibited strong *in vitro* antagonism against the major mycoparasites responsible for cobweb, dry bubble, wet bubble, and green mould diseases, consistent with findings for *B. velezensis* Kos (Clarke et al. 2022a; Clarke et al. 2022b). Genome sequencing revealed multiple biosynthetic gene clusters predicted to encode diverse non-ribosomal peptides and polyketide antimicrobials. However, targeted metabolite analyses identified fengycins as the dominant specialised metabolites produced under the culture conditions tested, suggesting these lipopeptides are the primary contributors to the observed antifungal and surfactant activities. Nonetheless, additional antimicrobial compounds may be produced only in response to specific cues, such as microbial competition or the casing environment, as many biosynthetic gene clusters remain silent under standard laboratory conditions and are activated only during co-culture or under particular environmental stimuli (Ochi 2017).

The chemical structures of the fengycin analogues produced by CM5, CM19 and CM35 were determined under *in vitro* culture conditions. Despite their high genomic similarity, CM19 yielded markedly higher levels of the fengycin analogue, suggesting that epigenetic regulation or one or more SNPs identified in these strains (Table S10) may influence lipopeptide production.

Fengycin exerts antifungal activity by forming pore-like complexes in fungal membranes in a lipid-dependent manner (Zakharova et al. 2019). The observation that fengycins are the major antifungal metabolites produced *in vitro*, combined with stronger antagonism toward mycoparasites (Hypocreales, Ascomycota) than toward the host, *A. bisporus* (Agaricales, Basidiomycota), suggests that differential membrane properties between these fungal groups may underlie this selective activity. This highlights the potential of fengycin-producing strains and engineered variants designed to minimise production of broad-spectrum antifungals, as targeted biocontrol agents in mushroom cultivation. It further underscores the promise of fengycins, and other lipid-specific membrane-active compounds, as selective fungicides. Ongoing work aims to optimise large-scale fengycin fermentation (Yin et al. 2024), and future research should characterise the activity, efficiency potential and selectivity of different fengycin analogues to guide industrial production.

Although fengycin is the most plausible contributor to the *in vitro* antifungal activity observed, this observation could not yet be experimentally verified through targeted mutagenesis, as the *B. velezensis* strains studied were highly recalcitrant to genetic transformation. The low but successful transformation of CM19 with pHAPII-gfp indicates that optimisation of transformation protocols may eventually enable disruption of the fengycin biosynthetic cluster. Additionally, neither expression of fengycin biosynthetic genes nor fengycin production could be detected when the strains were applied to casing (data not shown), consistent with their rapid persistence decline and the lack of disease suppression against *L. fungicola* in crop trials. Thus, to fully exploit fengycin-producing bacteria as biocontrol agents, future work will need to focus either on developing fengycins themselves as selective fungicides or on optimising bacterial formulation, survival, and metabolite production within the crop environment.

## Abbreviations

NGS: next generation sequencing
CD: cobweb disease
DBD: dry bubble disease
WBD: wet bubble disease
PLFA: phospholipid fatty acid

## 8. Declarations

### Ethical Approval

Not applicable

### Availability of data and materials

All data not already found in the manuscript is available on request from the corresponding author.

## 10. Author Contributions

Conceptualization: JC WK SK GP

Data curation: WK JC SK MK MJC TB

Formal analysis: JC MLT MT AT GP

Funding acquisition: JC SK MSRC JVW GP

Investigation: WK JC SK MK MJC TB MSRC JVW TB GP

Methodology: WK JC SK MK MJC

Project administration: JC SK MSRC JVW GP

Resources: WK JC SK MK MJC TB MSRC JVW TB GP

Software: WK JC SK MK MJC TB MSRC JVW TB GP

Supervision: JC SK MSRC JVW GP

Validation: WK JC SK MK MJC TB MSRC JVW TB GP

Visualization: WK JC GP

Writing – original draft: WK JC GP

Writing – review & editing: WK JC SK MK MJC TB MSRC JVW TB GP

## 11. Acknowledgements

*B. velezensis* CM5, CM19 and CM35 are protected as biocontrol agents against mushroom parasites in a patent published as EP4200400A1; WO2021255181A1PCT. 2023/06/28. The project leading to this report has received funding from the European Union’s Horizon 2020 research and innovation programme GA: 101000651 (BIOSCHAMP) and the Marie Sklodowska-Curie IF GA: 742966 (MYCOBIOME). JC is the recipient of a Ramon y Cajal contract [RYC2021-032796-I], funded by MCIN/AEI/10.13039/ 501100011033 and the European Union “NextGenerationEU”/PRTR”. We would also like to thank all members of the BIOSCHAMP consortium for all their helpful feedback.

