## Supplemental for "Biocontrol of mushroom crop mycoparasites by novel *Bacillus velezensis* strains"

### Supplementary Information

#### Supplementary Methods

**Methods S1: Specificity of the TaqMan assays:** The specificity of the assays was tested using genomic DNA of 27 strains. For most strains 0.5 ng of purified DNA was used, but for some strains a suspension of $10^9$- $10^{10}$ cells/mL was boiled for 10 min in 150 µL MilliQ water (MQ) prior to testing. For each TaqMan assay 2 µL of DNA or the boiled suspensions was mixed with 2 µL reaction mix containing PerfeCTa multiplex qPCR ToughMix 5x (Quantabio, Beverly, USA), 100 nM probe and 300 nM of each forward and reverse primer. The reactions were performed in a 12K Flex QuantStudio or a QuantStudio 5 Real-Time PCR system (Applied biosystems) using the following conditions: 95 °C for 2 min; 40 cycles of 95 °C for 15 s followed by 60 °C for 60 s. Analysis of the data was done by automatic threshold calculation within the Applied Biosystems software. A Ct value ≤ 35 was considered positive.

The specificity and usability were further tested using constituents of casing soils or casing soils of various compositions. DNA was extracted from 10 g of material using the DNeasy PowerMax Soil kit (Qiagen, Germany) according to the manufacturer's protocol. TaqMan assays were performed as described above with 2 µL DNA.

**Methods S2: Triplex TaqMan design:** The two simplex assays for the target bacteria were combined with an assay that quantifies *Xanthomonas campestris* pv. *campestris* (Xcc) into a triplex TaqMan (Köhl et al. 2011; Taparia et al. 2020). For each group of biostimulants, a triplex TaqMan assay was designed with two target-specific assays, each with its own dye (Cy5 or FAM) and an amplification and extraction control (*X. campestris*), labelled with HEX. The triplex TaqMan was performed using the same PCR conditions and materials as the simplex tests, with 100 nM probe and 300 nM of each primer in a total volume of 25 µL.

**Methods S3: Limit of detection of the assays:** The limit of detection for the triplex assays designed for *B. velezensis* strains CM19, CM5 and CM35 (Table S7) was determined in two casing soils, i.e. in peat and steamed spent casing soil. 100 µL of a ten-fold serial dilution of 3-days old cultures of the strains in Ringers in a PowerBead Pro tube was mixed with 250 mg of the casings ($4 \times 10^7 – 4 \times 10^1$ cells $g^{-1}$ of casing soil). Casing soils with 100 µL Ringers without supplemented bacteria served as a control. A volume of 100 µL of a suspension of *X. campestris* (Xcc) of $10^6$ cell $mL^{-1}$ was added to each sample as extraction and amplification control. The soils were subsequently freeze dried overnight and shaken for 2 times 90 seconds in a paint shaker. DNA was extracted using a DNeasy Power Soil Pro Kit according to the manufacturer's instructions. The triplex TaqMan assay was executed as described above with 0.75 µL of each primer (10 µM) and 0.75 µL of each probe (10 µM), labelled with FAM, Cy5 (for target specific probes) or HEX (for Xcc).

### Supplementary Tables

**Table S1: Media employed to isolate and culture bacteria from the casing and *A. bisporus* basidiomes.**

| Media | Recipe |
|---|---|
| Agar-Casing-LBA | 150 g of dried and milled fresh casing are autoclaved with approx. 800 mL of distilled water at 121ºC and 1 at. The supernatant is filtered through a double layer of muslin to remove the bulk of the substrate, and 10 g of tryptone (Cultimed Panreac #403682, Barcelona, Spain), 5 g of yeast extract (Cultimed #A3732), + 10 g NaCl (Thermo Scientific #207790010) and 15 g of bacteriological agar (Cultimed #402302) are added to the supernatant liquid up to 1 L with distilled water. |
| Agar-Compost | 50 g of dried and ground phase II compost for *A. bisporus* are autoclaved with approx. 800 mL of distilled water at 121ºC and 1 at. The supernatant is filtered through a double layer of muslin to remove the bulk of the substrate, and 15 g of bacteriological agar (Cultimed #402302) is added up to 1 L with distilled water. |
| King´s agar + CFC | Selective medium for *Pseudomonas*: Peptone-protease No.3 (Thermo Scientific #211693), 20 g $L^{-1}$; $K_2HPO_4$ (Thermo Scientific #215472500), 1.5 g $L^{-1}$; $MgSO_4$ (Thermo Scientific #A14491.0I), 1.5 g $L^{-1}$; Bacteriological Agar, 15 g $L^{-1}$ (Cultimed #402302); Glycerol (Thermo Scientific #A16205.AP), 10 mL $L^{-1}$, plus CFC Pseudomonas supplement (Thermo Scientific #SR0103E) (content of a vial for 500 mL of medium): cetrimide (5 mg); Fusionic acid (5 mg); Cephalosporin (25 mg). |
| LBA | 10 Tryptone (Cultimed, Barcelona, Spain) + 5 g Yeast extract (Cultimed) + 10 g NaCl (Thermo Scientific) + 15 g Bacteriological Agar (Cultimed) for 1L; |
| TSA | Tryptone soya agar was prepared according to the manufacturer's instructions (Cultimed #416106) |
| HiCrome™ B. Agar | Selective medium for *B.* was prepared according to the manufacturer's instructions (HiMedia Labs, India). |
| PBS electroporation buffer | PEB: 272 mM sucrose, 1 mM $MgCl_2 \cdot 6H_2O$, 7 mM phosphate buffer, pH 7.4) |

**Table S2: Cultivable bacterial microbiome from the peat casing layer and basidiomes.**

| Strain code | Origin (isolated from)[1] | Strain (based on sequencing of 16S rRNA)[2] |
|---|---|---|
| CM5 | CM | *Bacillus velezensis* (HQ831396.1) - 98.76% |
| CM19 | CM | *B. velezensis* (MT184842.1) - 83.25% |
| CM35 | CM | *B. velezensis* (MF581451.1) - 99.43% |
| B10 | B | *Pantoea* sp. (MK398041.1) - 82.66% |
| B13 | B | *Serratia* sp. (MT623418.1) - 98.21% |
| B18 | B | *Staphylococcus fleurettii* (MN758802.1) - 98.43% |

| | | |
|---|---|---|
| B19 | B | *Staphylococcus* sp. (KT029134.1) - 99.25% |
| B9A | B | *Serratia* sp. (KF791514.1) - 98.10% |
| B9B | B | *Vibrio ruber* (MT422024.1) - 84.38% |
| B9C | B | *Pseudochrobactrum asaccharolyticum* (KF263562.1) - 99.21% |
| B11W | B | *Staphylococcus fleurettii* (MK015775.1) - 99.30% |
| B11Y | B | *Arthrobacter* sp. (MG874752.1) - 99.35% |
| B14A | B | *Serratia marcescens* (MT645673.1) - 98.21% |
| B14B | B | *Paenochrobactrum* sp. (JF804769.1) - 98.92% |
| B15W | B | Staphylococcaceae bacterium (AY841364.1) - 77.00% |
| B15Y | B | *Glutamicibacter* sp. (MW627371.1) - 100% |
| B16 | B | *Pseudomonas* sp. (MK949377.1) - 98.70% |
| B17A | B | *Klebsiella oxytoca* (MF144451.1) - 98.90% |
| B17B | B | *Pseudochrobactrum* sp. (CP075349.1) - 82.13% |
| B20A | B | *Bacillus* sp. (KM242479.1) - 95.80% |
| B20B | B | Uncultured bacterium (DQ817818.1) - 85.82% |
| B21A | B | *Glutamicibacter mishrai* (CP032549.1) - 99% |
| B21B | B | *Staphylococcus* sp. (MT516468.1) - 92.47% |
| B21C | B | *Pseudomonas putida* 1616X1 -98% |
| B22A | B | *Pseudomonas putida* JQ581 92% |
| B22B | B | *Staphylococcus saprophyticus* poll1 - 96.69% |
| CM1 | CM | *Pseudomonas* sp. (JF430830.1) - 79.95% |
| CM2 | CM | *Pseudomonas* sp. (MN231751.1) - 98.25% |
| CM3 | CM | *Serratia* sp. strain AXJ-M - 98.62 |
| CM4 | CM | *Serratia* sp. (MN928759.1) - 94.85% |
| CM6 | CM | *Serratia* sp. (MT472080.1) - 94.93% |
| CM7 | CM | *Serratia* sp. (MT472080.1) - 96.18% |
| CM10 | CM | *Serratia* sp. (MT472080.1) - 94.21% |
| CM11 | CM | *Bacillus* sp. (DQ365574.1) - 78.31% |
| CM12 | CM | *Chryseobacterium* sp. (KX928035.1) - 80.14% |
| CM13 | CM | No significant similarity found |
| CM14 | CM | *Pseudomonas* sp. (KF263605.1) - 91.85% |
| CM15 | CM | *Brevundimonas* sp. (MK110476.1) |
| CM16 | CM | Uncultured bacterium (KX507468.1) - 73.16% |
| CM17 | CM | *Comamomas* sp. (KC294078.1) - 99.33% |

| | | |
|---|---|---|
| CM18 | CM | *B. velezensis* LA-33 - 95% |
| CM20 | CM | *B. velezensis* WC-14 - 95.15% |
| CM21 | CM | *B. velezensis* YJ11-1-4 = 90.71% |
| CM22 | CM | *Pseudomonas putida* (MN795707.1) - 88.42% |
| CM23 | CM | *B. licheniformis* (MN756663.1) - 97.04% |
| CM25 | CM | *Pseudomonas* sp. (JX678920.1) - 78.40% |
| CM26 | CM | *Bacillus* sp. (MN989043.1) - 98.43% |
| CM27 | CM | *Acidovorax radicis* IMCC34918 - 94.59% |
| CM30 | CM | *Pseudomonas* sp. (MT568555.1) - 98.55% |
| CM31 | CM | *Rhodococcus* sp. CP5 - 86.34% |
| CM32 | CM | *Glutamicibacter halophytocola* (MT533909.1) - 96.24% |
| CM33 | CM | *Pseudomonas* sp. (KF202578.1) - 78.63% |
| CM34 | CM | *Serratia* sp. (MT472080.1) - 95.33% |
| CM36 | CM | *Serratia* sp. strain TC/03 - 96.44% |
| CM37 | CM | *Serratia* sp. (MT472080.1) - 95.29% |
| CM38 | CM | *Klebsiella* sp. (MK209044.1) - 87.85% |
| CM39 | CM | *Pseudomonas frederiksbergensis* (MG561792.1) - 74.47% |
| CM40 | CM | *Pseudomonas* sp. - Uncultured bacterium (GQ101186.1) - 74.39% |
| CM41 | CM | *Pseudochrobactrum* sp. (MK095772.1) - 81.12% |
| CM42 | CM | *Stenotrophomonas* sp. (MT258986.1) - 84.93% |
| CM43 | CM | *Glutamicibacter* sp. (MK841094.1) - 97.99% |
| EM1 | B | *Bacillus algicola* (MN746261.1) - 96.45% |
| EM2 | B | *Bacillus* sp. (FN646629.1) - 97.80% |
| EM3 | B | *Escherichia coli* (CP044298.1) - 89.26% |
| EM4 | B | *Kluyvera intermedia* (MT081698.1) - 93.37% |
| EM5 | B | *B. velezensis* (MN272328.1) - 99.28% |
| EM6 | B | *Staphylococcus succinus* (KX348384.1) - 98.62% |
| EM7 | B | *Bacillus firmus* (KY316472.1) - 99.33% |
| EM8 | B | *Bacillus licheniformis* (JX041900.1) - 75.69% |
| EM9 | B | Uncultured *Pseudomonas* sp. (HQ616313.1) - 79.01% |
| EM10 | B | *Stenotrophomonas* sp. (KC853134.1) - 83.03% |
| EM11 | B | *Pseudomonas fluorescens* (JX484838.1) - 84.65% |
| EM12 | B | *Pseudomonas* sp. P.f-9103 (KM032182.1) - 90.31% |
| EM15S | B | *Stenotrophomonas rhizophila* (JQ670922.1) - 89.31% |

| | | |
|---|---|---|
| EM15L | B | *Rhodococcus* sp. (MH043947.1) - 95.93% |
| EM16 | B | *Lampropedia* sp. BRTC-3 (KU612256.1) - 97.10% |
| EM17 | B | *Rhodococcus* sp. (JF742932.1) - 75.57% |
| EM18 | B | Enterobacteriaceae bacterium (AB275371.1) - 79.66% |
| EM19 | B | *Pseudomonas* sp. SB11-07 (GU595316.1) - 82.58% |
| EM21 | B | *Pseudomonas* sp. (KX264924.1) - 98.47% |
| EM22 | B | *Pseudomonas* sp. (JN201547.1) - 98.19% |
| EM23 | B | *Pseudomonas* sp. (KU510013.1) - 98.67% |
| EM24 | B | *Ochrobactrum* sp. (MT271895.1) - 99.70% |
| EM25 | B | *Pseudomonas* sp. (MW737403.1) - 98.11% |
| EM27 | B | *Paracoccus* sp. (KY971629.1) - 99.38% |
| EM28 | B | *Stenotrophomonas maltophilia* (DQ339644.1) - 90.60% |
| EM29 | B | *Pseudomonas aeruginosa* (CP042269.1) - 92.67% |
| EM30 | B | *Microbacterium* sp. (MT576542.1) - 98.51% |
| EM31 | B | *Microbacterium* sp. (JQ977247.1) - 97.67% |
| EM32 | B | *Serratia* sp. (AM231089.1) - 89.77% |
| EM33 | B | *Rahnella inusitata* (MT555345.1)- 91.28% |
| EM34 | B | *Bacillus* sp. (GQ199751.1) - 96.04% |
| EM35 | B | *Staphylococcus* sp. (KR029198.1) - 97.27% |
| EM36 | B | Uncultured bacterium-Enterobacteria (KP142371.1) - 72.48% |
| EM37 | B | *Staphylococcus* sp. (MN103949.1) - 98.85% |
| EM38 | B | *Microbacterium* sp. (MH130312.1) - 92.11% |
| EM39 | B | *Bacillus amyloliquefaciens* (KJ639041.1) - 84.50% |
| EM41 | B | *Citrobacter* sp. SI-GU-1B (KP322786.1) - 78.91% |
| EM42 | B | *Pseudomonas fluorescens* (FJ787327.1) - 84.16% |
| EM43 | B | *Chryseobacterium* sp. (DQ530065.1) - 91.17% |
| EM44 | B | *Pseudomonas* sp. (MK371440.1) - 83.87% |
| EM45 | B | *Pseudomonas* sp.185 - 97.62% |
| EM46 | B | *Pseudomonas koreensis* (MW405680.1) -88.78% |
| EM47 | B | *Pseudomonas* sp. (MW433970.1) - 97.37% |
| EM48 | B | *Pseudomonas mosselii* (KY426038.1) - 97.48% |
| EM49 | B | *Exiguobacterium* sp. (MT611261.1) - 97.89% |
| EM13 | B | *Chryseobacterium* sp. (KX079844.1) - 91.73% |
| EM14 | B | *Enterobacter* sp. (FJ587228.1) - 79.77% |

| | | |
|---|---|---|
| EM20 | B | *Chryseobacterium* sp. (KX079844.1) - 87.50% |
| EM26 | B | *Chryseobacterium* sp. (MG893574.1) - 90.68% |
| EM40 | B | *Serratia marcescens* (EF635971.1) - 89.23% |
| CM8 | CM | *Pseudomonas laurentiana* SeaQual_P_B70/W - 89% |
| CM9 | CM | *Pseudomonas putida* (AF307869.1) - 99.16% |
| CM24 | CM | *Pseudomonas* sp. (KM391420.1) - 96.38% |
| CM28 | CM | Uncultured *Pseudomonas* sp. (EU029490.1) - 82.52% |
| CM29 | CM | *Serratia grimesii* (DQ481467.1) - 84.32% |
| CM44 | CM | Uncultured *Pseudomonas* sp. (HM011771.1) - 88.41% |
| CM45 | CM | *Glutamicibacter* sp. (MK847918.1) - 98.04% |
| CM46 | CM | *Citrobacter werkmanii* (LR699014.1) - 83.62% |
| CM47 | CM | *Ochrobactrum* sp. (KY678891.1) - 89.09% |
| CM48 | CM | *Micrococcus endophyticus* (MT214326.1) - 100% |
| CM49 | CM | *Streptomyces* sp. (KF650632.1) - 76.03% |
| CM50 | CM | *Pseudochrobactrum* sp. (KF937789.1) - 83.43% |
| CM51 | CM | *Citrobacter freundii* (MW940831.1) - 79.31% |
| CM52 | CM | Alcaligenaceae bacterium (MH593839.1) - 89.41% |
| CM53 | CM | *Rhodococcus* sp. (MT631993.1) - 100% |
| CM54 | CM | *Brevundimonas* sp. (MF111489.1) - 96.57% |
| CM56 | CM | *Pseudomonas plecoglossicida* (KF261039.1) - 100% |
| CM57 | CM | *Pseudomonas* sp. WB4.4-63 (AM934697.1) - 84.82% |
| CM58 | CM | No significant similarity found |
| CM59 | CM | Uncultured bacterium (KP156579.1) - 75.83% |
| CM60 | CM | *Brevundimonas diminuta* (KC152996.1) - 75.27% |
| CM61 | CM | *Brevundimonas staleyi* EB109 - 88.22% |
| CM62 | CM | *Enterobacter* sp. (MT557025.1) - 75.00% |
| CM63 | CM | *Bacillus oceanisediminis* SH89 - 91.70% |
| CM55 | CM | *Pseudomonas paralactis* BF388 - 94.97% |

**[1]The strains were isolated either from casing material (CM) or adult basidiomes (B). [2]BLASTn (blast.ncbi.nlm.nih.gov) was used to classify the microbes based on their 16s rRNA sequences.**

**Table S3: Quality Assessment of *B. velezensis* genomes using QUAST.**

| Statistics without reference[1] | 064048E_CM5 | 064049E_CM35 | 064050E_EM5 | 064051E_EM39 |
|---|---|---|---|---|
| # contigs | 17 | 16 | 17 | 11 |
| # contigs (>= 0 bp) | 46 | 32 | 32 | 48 |

| | | | | |
|---|---|---|---|---|
| # contigs (>= 1000 bp) | 13 | 13 | 14 | 10 |
| # contigs (>= 5000 bp) | 8 | 8 | 9 | 7 |
| # contigs (>= 10000 bp) | 8 | 8 | 9 | 7 |
| # contigs (>= 25000 bp) | 7 | 7 | 8 | 6 |
| # contigs (>= 50000 bp) | 7 | 7 | 8 | 6 |
| Largest contig | 2233836 | 2233835 | 2233835 | 2061313 |
| Total length | 4203474 | 4202775 | 4202331 | 3891334 |
| Total length (>= 0 bp) | 4213899 | 4207978 | 4206541 | 3902642 |
| Total length (>= 1000 bp) | 4200517 | 4200358 | 4199939 | 3890670 |
| Total length (>= 5000 bp) | 4192636 | 4192415 | 4192079 | 3885132 |
| Total length (>= 10000 bp) | 4192636 | 4192415 | 4192079 | 3885132 |
| Total length (>= 25000 bp) | 4175744 | 4175523 | 4175187 | 3868234 |
| Total length (>= 50000 bp) | 4175744 | 4175523 | 4175187 | 3868234 |
| N50 | 2233836 | 2233835 | 2233835 | 2061313 |
| N75 | 488218 | 488218 | 488307 | 1012020 |
| L50 | 1 | 1 | 1 | 1 |
| L75 | 3 | 3 | 3 | 2 |
| GC (%) | 45.84 | 45.85 | 45.84 | 46.43 |
| Mismatches | | | | |
| # N's | 0 | 0 | 0 | 0 |
| # N's per 100 kbp | 0 | 0 | 0 | 0 |

**[1]All statistics are based on contigs of size >= 500 bp, unless otherwise noted (e.g., "# contigs (>= 0 bp)" and "Total length (>= 0 bp)" include all contig**

**Table S4: Primers used in this study for the phylogenetic analysis of isolated *B. velezensis* strains.**

| Primer name | Sequence (5' - 3') | Ref. |
|---|---|---|
| 16s_27F | AGAGTTTGATCCTGGCTCAG | Taguchi *et al.*, 2008 |
| 16s_1492R | GGTTACCTTGTTACGACTT | Taguchi *et al.*, 2008 |
| rpoB24F | CGCATGATTTGAGGGG | Fajardo-Cavazos *et al.*, 2018 |
| rpoB737R | GGCGGCTCTCCAGG | Fajardo-Cavazos *et al.*, 2018 |
| tufGPF | ACGTTGACTGCCCAGGACAC | Draganic *et al.*, 2017 |
| tufGPR | GATACCAGTTACGTCAGTTGTACGGA | Draganic *et al.*, 2017 |
| WK_rpoD_F | CCACGAAACAGAAACAGAACTTAC | This paper |
| WK_rpoD_R | CTGCTCGGATGTCTCAATTTG | This paper |
| WK_pgk_2_F | CATCGACGTAAAAGGCAAAG | This paper |

| | | |
|---|---|---|
| WK_pgk_2_R | CTACACCTGGAAGCTCTTTG | This paper |
| Wk_gyrB_2_F | GTATTAGAAGGTTTGGAAGCTGTTCG | This paper |
| WK_gyrB_2_R | CATTTCTCCAAGACCTTTATAACGC | This paper |

**Table S5: Experimental blocks used in crop trial**

| Treatment | Code | Dose[1] | Infected (*L. fungicola* 150/1)[2] |
|---|---|---|---|
| **Control** | Control+ | 20 ml Tap water | $10^6$ conidia $m^{-2}$ |
| **Control** | Control- | 20 ml Tap water | - |
| ***B. velezensis* CM5** | CM5 | $10^9$ cfu $m^{-2}$ | $10^6$ conidia $m^{-2}$ |
| ***B. velezensis* CM19** | CM19 | $10^9$ cfu $m^{-2}$ | $10^6$ conidia $m^{-2}$ |
| ***B. velezensis* CM35** | CM35 | $10^9$ cfu $m^{-2}$ | $10^6$ conidia $m^{-2}$ |
| **Prochloraz-Mn** | PCL | 1 g $m^{-2}$ | $10^6$ conidia $m^{-2}$ |

**[1]PCL was applied 4 days after casing, bacteria were applied 5 days after casing. cfu: colony forming unit.**
**[2]Conidia were applied 10 days after casing.**

**Table S6. Specificity of TaqMan assays developed for *B. velezensis* strains selected as biocontrol agents for *A. bisporus* crops.**

| Species | [1]Strain No | Geographic origin | Yr of isolation | Isolated from: | [2]*B. velezensis* TaqMan (Ct) | |
|---|---|---|---|---|---|---|
| | | | | | 2HP2.21 | HP2.19 |
| *Pseudomonas putida* | PMS118R | New Zealand | 1990 | Casing - mushroom farms | ND | ND |
| *Pseudomonas fluorescens* | SBW25 | United Kingdom | 1989 | *Beta vulgaris* | ND | ND |
| *P. fluorescens* | PMS232 | United Kingdom | 1989 | *Beta vulgaris* | ND | ND |
| *P. fluorescens* | 55 | | | | ND | ND |
| *Pseudomonas* sp. | B18 | Spain | 2017 | Casing - mushroom farms (this study) | ND | ND |
| *Pantoea* sp. | B11 | Spain | 2017 | Casing - mushroom farms (this study) | ND | ND |
| *Serratia* sp. | B4 | Spain | 2017 | Casing - mushroom farms (this study) | ND | ND |
| *B. velezensis* | CM19 | Spain | 2017 | Casing - mushroom farms (this study) | 24.2 | 23.4 |
| *B. velezensis* | CM5 | Spain | 2017 | Casing - mushroom farms (this study) | 24.2 | 24 |

| | | | | | | |
|---|---|---|---|---|---|---|
| *B. velezensis* | CM35 | Spain | 2017 | Casing - mushroom farms (this study) | 23.7 | 23.1 |
| *Bacillus sp.* | B20A | Spain | 2017 | Casing - mushroom farms (this study) | ND | ND |
| *Bacillus sp.* | CM11 | Spain | 2017 | Casing - mushroom farms (this study) | ND | ND |
| *B. velezensis* | CM18 | Spain | 2017 | Casing - mushroom farms (this study) | 15 | 14.1 |
| *B. velezensis* | CM20 | Spain | 2017 | Casing - mushroom farms (this study) | 15.8 | 14.9 |
| *B. velezensis* | CM21 | Spain | 2017 | Casing - mushroom farms (this study) | 15.5 | 14.1 |
| *Bacillus sp.* | CM26 | Spain | 2017 | Casing - mushroom farms (this study) | ND | ND |
| *Bacillus algicola* | EM1 | Spain | 2020 | Mushroom endophyte (this study) | ND | ND |
| *Bacillus sp.* | EM2 | Spain | 2020 | Mushroom endophyte (this study) | ND | ND |
| *B. velezensis* | EM5 | Spain | 2020 | Mushroom endophyte (this study) | 15.1 | 14.3 |
| *Bacillus firmus* | EM7 | Spain | 2020 | Mushroom endophyte (this study) | ND | ND |
| *Bacillus licheniformis* | EM8 | Spain | 2020 | Mushroom endophyte (this study) | ND | ND |
| *Bacillus sp.* | EM34 | Spain | 2020 | Mushroom endophyte (this study) | ND | ND |
| *B. velezensis* | EM39 | Spain | 2020 | Mushroom endophyte (this study) | ND | ND |
| *B. subtilis* | KDSA | - | - | - | ND | ND |
| *B. thuringiensis* | 407 cry- | France | 198? | Isolated as a lepidopteran-active strain (acrystalliferous derivative) | ND | ND |

| | | | | | | |
|---|---|---|---|---|---|---|
| *B. subtilis* | 168 | USA | 194? | Model organism | ND | ND |
| | [3]MQ | | | | ND | ND |

**[1] Target strains are highlighted in green cells. [2] Per strain, two TaqMan assays were used (2HP2.21 and HP2.19). All probes were labelled with FAM (6-carboxyfluorescein, green-fluorescent dye). ND = not detected after 40 cycles. Ct (Cycle threshold): value is the number of PCR cycles required for the fluorescent signal to cross a certain threshold, indicating detectable levels of the target nucleic acid. [3]MQ= Molecular Quantification.**

**Table S7: TaqMan primers and probes for *B.* velezensis (strains CM5, CM19; CM35) and for the amplification and extraction control *X. campestris* pv. *campestris*.**

| | **Species** | ***B. velezensis*** | | ***X. campestris* pv. *campestris*** |
|---|---|---|---|---|
| | **Strain No.** | **CM19; CM5; CM35** | | **Extraction Control** |
| | **Target gene** | **Hypothetical protein** | **Hypothetical protein** | **Unknown** |
| **Forward primer** | Name | *B. velezensis*_HP2.21_set2_F | *B. velezensis*_HP2.19_set3_F | XccF |
| | Sequence (5'-3') | CCCTGAACATAG GGTGTATGATG | CATTATTTCAGACAC AGATGCTCTT | GTGCATAGGCCAC GATGTTG |
| **Reverse Primer** | Name | *B. velezensis* _HP2.21_set2_R | *B. velezensis* _HP2.19_set3_R | XccR |
| | Sequence (5'-3') | ACAGCAATTGGT TGGAAGAAAC | TGAAGACATTATTGA ATGCTTACCC | CGGATGCAGAGCG TCTTACA |
| **Probe** | Name | *B. velezensis* _HP2.21_P | *B. velezensis* _HP2.19_P | XccP |
| | Sequence (5'-3') | TTTGGAATCCCAC GTTCGTTTACA | ACCTGCTGACTGTTT GCAAACTCT | CAAGCGATGTACT GCGGCCGTC |
| | Dye[1] | FAM | Dy5 | HEX |

**[1]Label in Triplex assay.**

**Table S8: Results of antimicrobial confrontation assays of cultivable bacterial microbiome from the peat casing layer and positive control strains NZ011, NZI7 and NCPPB2192 against 4 major fungal parasites of *A. bisporus*: (i) green mould/*Trichoderma* aggressivum TAV1 (Trich), (ii) wet bubble/*Mycogone perniciosa* M25 (Myco), (iii) dry bubble/*L. fungicola* 150/1 (Lec), and (iv) cobweb/*Cladobotryum mycophilum* CM13900 (Clad).**

| **Code** | **Strain (Seq. result 16S rRNA)** | **Trich** | **Clad** | **Myco** | **Lec** |
|---|---|---|---|---|---|
| CM5 | *B. velezensis* CM5 | *1 | 1 | 1 | 1 |
| CM19 | *B. velezensis* CM19 | 1 | 1 | 1 | 1 |
| CM35 | *B. velezensis* CM35 | 1 | 1 | 1 | 1 |
| NZ011 | *Pseudomonas fluorescens* NZ011 | 1 | 1 | 1 | 1 |
| NZI7 | *Pseudomonas fluorescens* NZI7 | 1 | 1 | 1 | 1 |
| NCPPB2192 | *Pseudomonas tolasasii* NCPPB 2192 (smooth) | 1 | 1 | 0 | 0 |
| B16 | *Pseudomonas* sp. (MK949377.1) - 98.70% | 0 | 0 | 0 | 0 |
| CM1 | *Pseudomonas* sp. (JF430830.1) - 79.95% | 0 | 0 | 0 | 0 |

| | | | | | |
|---|---|---|---|---|---|
| CM2 | *Pseudomonas* sp. (MN231751.1) - 98.25% | 0 | 0 | 0 | 0 |
| CM11 | *Bacillus* sp. (DQ365574.1) - 78.31% | 0 | 0 | 0 | 0 |
| CM14 | *Pseudomonas* sp. (KF263605.1) - 91.85% | 0 | 0 | 0 | 0 |
| CM18 | *B. velezensis* strain LA-33 - 95% | 1 | 1 | 1 | 1 |
| CM20 | *B. velezensis* strain WC-14 - 95.15% | 1 | 1 | 1 | 1 |
| CM21 | *B. velezensis* strain YJ11-1-4 = 90.71% | 1 | 1 | 1 | 1 |
| CM22 | *Pseudomonas putida* (MN795707.1) - 88.42% | 0 | 0 | 0.75 | 0 |
| CM23 | *Bacillus licheniformis* (MN756663.1) - 97.04% | 0 | 0 | 0 | 0 |
| CM25 | *Pseudomonas* sp. (JX678920.1) - 78.40% | 0 | 0 | 0 | 0 |
| CM26 | *Bacillus oceanisediminis* strain Xmb066 - 98.47% | 0 | 0 | 0 | 0 |
| CM30 | *Pseudomonas* sp. ( MT568555.1) - 98.55% | 0 | 0 | 0 | 0 |
| CM33 | *Pseudomonas* sp. (KF202578.1) - 78.63% | 1 | 1 | 1 | 0 |
| CM39 | *Pseudomonas frederiksbergensis* (MG561792.1) 74.47% | 0 | 0 | 1 | 0 |
| CM40 | *Pseudomonas* sp. Uncultured (GQ101186.1) - 74.39% | 0 | 1 | 0 | 0 |
| EM5 | *B. velezensis* (MN272328.1) - 99.28% | 1 | 1 | 1 | 1 |
| EM9 | Uncultured *Pseudomonas* sp. (HQ616313.1) - 79.01% | 0 | 0 | 0 | 0 |
| EM11 | *Pseudomonas fluorescens* (JX484838.1) - 84.65% | 0 | 0 | 1 | 0 |
| EM12 | *Pseudomonas* sp. P.f-9103 (KM032182.1) - 90.31% | 0 | 0 | 0 | 0 |
| EM19 | *Pseudomonas* sp. SB11-07 (GU595316.1) - 82.58% | 0 | 0 | 1 | 0 |
| EM21 | *Pseudomonas* sp. (KX264924.1) - 98.47% | 0 | 0 | 0.33 | 0 |
| EM22 | *Pseudomonas* sp. (JN201547.1) - 98.19% | 0 | 0 | 0 | 0 |
| EM23 | *Pseudomonas* sp. (KU510013.1) - 98.67% | 0 | 0 | 0 | 0 |
| EM25 | *Pseudomonas* sp. (MW737403.1) - 98.11% | 0 | 0 | 0 | 0 |
| EM29 | *Pseudomonas aeruginosa* (CP042269.1) - 92.67% | 0 | 0 | 0 | 0 |
| EM39 | *Bacillus amyloliquefaciens* (KJ639041.1) - 84.50% | 1 | 1 | 0.75 | 1 |
| EM42 | *Pseudomonas fluorescens* (FJ787327.1) - 84.16% | 0 | 0 | 0 | 0 |
| EM44 | *Pseudomonas* sp. (KX264924.1) - 98.47% | 0 | 0 | 0 | 0 |
| EM46 | *Pseudomonas koreensis* (MW405680.1) -88.78% | 1 | 1 | 1 | 0 |
| EM47 | *Pseudomonas* sp. (MW433970.1) - 97.37% | 0 | 0 | 0 | 0 |
| EM48 | *Pseudomonas mosselii* (KY426038.1) - 97.48% | 1 | 1 | 1 | 0 |
| CM9 | *Pseudomonas putida* (AF307869.1) - 99.16% | 0 | 0 | 0 | 0 |
| CM24 | *Pseudomonas* sp. (KM391420.1) - 96.38% | 0 | 0 | 0 | 0 |
| CM28 | Uncultured *Pseudomonas* sp. (EU029490.1) - 82.52% | 1 | 1 | 1 | 0 |
| CM44 | Uncultured *Pseudomonas* sp. (HM011771.1) - 88.41% | 0 | 0 | 0 | 0 |

| | | | | | |
|---|---|---|---|---|---|
| CM56 | *Pseudomonas plecoglossicida* (KF261039.1) - 100% | | 1 | 1 | 0 |
| CM57 | *Pseudomonas* sp. WB4.4-63 (AM934697.1) - 84.82% | 0 | 0 | 0 | 0 |
| CM64 | *B. velezensis* QST 713 | 1 | 1 | 1 | 1 |
| CM32 | *Glutamicibacter halophytocola* (MT533909.1) - 96.24% | 0 | 0 | 0 | 0 |
| B21A | *Glutamicibacter mishrai* (CP032549.1) - 99% | 0 | 0 | 0 | 0 |
| CM43 | *Glutamicibacter* sp. (MK841094.1) - 97.99% | 0 | 0 | 0 | 0 |
| CM45 | *Glutamicibacter* sp. (MK847918.1) - 98.04% | 0 | 0 | 0 | 0 |
| B15Y | *Glutamicibacter* sp. (MW627371.1) - 100% | 0 | 0 | 0 | 0 |
| EM1 | *Bacillus algicola* (MN746261.1) - 96.45% | 0 | 0 | 0 | 0 |
| EM7 | *Bacillus firmus* (KY316472.1) - 99.33% | 0 | 0 | 0 | 0 |
| EM8 | *B. licheniformis* (JX041900.1) - 75.69% | 0 | 0 | 0 | 0 |
| EM2 | *Bacillus* sp. (FN646629.1) - 97.80% | 0 | 0 | 0 | 0 |
| EM34 | *Bacillus* sp. (GQ199751.1) - 96.04% | 0 | 0 | 0 | 0 |
| B20A | *Bacillus* sp. (KM242479.1) - 95.80% | 0 | 0 | 0 | 0 |

***Score: 1 – Antimicrobial activity observed when confronted to mycoparasites; 0 – No antimicrobial activity observed.**

**Table S9: Summary of genomic data for *B.* velezensis casing isolates compared to QST713 (BLASTn (Nucleotide BLAST); BLASTp (Protein-Protein BLAST).**

| | QST713 | CM5 | CM19 | CM35 | EM5 | EM39 |
|---|---|---|---|---|---|---|
| **Similarity to [1]QST713 (%)** | 100 | 99.99 | 99.99 | 99.99 | 99.99 | 98.38 |
| **Genes (total)** | 4,238 | 4,256 | 4,255 | 4,256 | 4,256 | 3,906 |
| **GC Content** | 46 | 45.84 | 46 | 45.85 | 45.84 | 46.43 |
| **Genome Size** | 4233757 | 4240819 | 4240819 | 4240818 | 4240818 | 3929792 |
| **[2]CDSs (total)** | 4,047 | 4,137 | 4,136 | 4,137 | 4,137 | 3,788 |
| **Genes (coding)** | 4,047 | 4,049 | 4,048 | 4,048 | 4,048 | 3,686 |
| **CDSs (with protein)** | 4,047 | 4,049 | 4,048 | 4,048 | 4,048 | 3,686 |
| **Genes (RNA)** | 107 | 119 | 119 | 119 | 119 | 118 |
| **[3]rRNAs (5S, 16S, 23S)** | 9, 8, 8 | 10, 9, 9 | 10, 9, 9 | 10, 9, 9 | 10, 9, 9 | 9, 9, 9 |
| **Complete rRNAs (5S, 16S, 23S)** | 9, 8, 8 | 10, 9, 9 | 10, 9, 9 | 10, 9, 9 | 10, 9, 9 | 9, 9, 9 |
| **[4]tRNAs** | 77 | 86 | 86 | 86 | 86 | 86 |
| **[5]ncRNAs** | 5 | 5 | 5 | 5 | 5 | 5 |
| **Pseudo Genes (total)** | 84 | 88 | 88 | 89 | 89 | 102 |
| **CDSs (without protein)** | 84 | 88 | 88 | 89 | 89 | 102 |
| **Pseudo Genes (ambiguous residues)** | 0 of 84 | 0 of 88 | 0 of 88 | 0 of 89 | 0 of 89 | 0 of 102 |

| | | | | | | |
|---|---|---|---|---|---|---|
| **Pseudo Genes (frameshifted)** | 48 of 84 | 51 of 88 | 51 of 88 | 52 of 89 | 52 of 89 | 61 of 102 |
| **Pseudo Genes (incomplete)** | 58 of 84 | 61 of 88 | 61 of 88 | 61 of 89 | 61 of 89 | 65 of 102 |
| **Pseudo Genes (internal stop)** | 14 of 84 | 11 of 88 | 11 of 88 | 11 of 89 | 11 of 89 | 9 of 102 |
| **Pseudo Genes (multiple problems)** | 30 of 84 | 29 of 88 | 29 of 88 | 29 of 89 | 29 of 89 | 30 of 102 |

**[1]Results for all strains are from analyses using NCBI PGAAP (Tatusova et al. 2016). Quality assessment of *B. velenzensis* genomes using QUAST can be found in Table S3; [2]CDS: Coding sequences; [3]rRNA: ribosomal RNA; [4]tRNA: transfer RNA; [5]ncRNA: non-coding RNA.**

**Table S10: Genes in novel *B.* velezensis strains containing SNPs which affect amino acid sequences compared to QST713.**

| Locus tag | Gene | Product | CM19 | CM35 | CM5 | EM5 | Amino acid differences[1] |
|---|---|---|---|---|---|---|---|
| BVQ_00080 | serS | Seryl-tRNA synthetase | 1 (0) | 1 (0) | 1 (0) | 1 (0) | Sub 188: E>K |
| BVQ_01790 | srfAA | Surfactin synthetase | 1 (0) | 1 (0) | 1 (0) | 1 (0) | No differences |
| BVQ_01795 | | | 27 (0) | 27 (0) | 27 (0) | 27 (0) | Sub 2822: V>I; Ins 2842-2842: E; Sub 2986: A>T; Sub 2989: A>S; Sub 3025: A>E |
| BVQ_04115 | | | 0 (2) | 0 (2) | 0 (2) | 0 (2) | Ins 1454: TGPTGSTGSTGETGTTGSTGATG. Del 1757: TGATGV |
| BVQ_04120 | | | 2 (0) | | | | Sub 413: V>I |
| BVQ_06715 | | | 21 (4) | 20 (4) | 20 (4) | 20 (4) | Sub 1014: V>I; Sub 1020: G>S; Sub 1024: S>E; Sub 1025: T>N; Sub 1028: G>T; Sub 1031: K>S |
| BVQ_07310 | | | | 0 (1) | | 0 (1) | Sub 133: P>T; Sub 134: K>E |
| BVQ_08470 | topA | DNA topoisomerase I | | 1 (0) | | | Sub 620: L>P |
| BVQ_09135 | nrdF | β subunit of class Ib ribonucleotide reductase (RNR) | 20 (0) | 20 (0) | 20 (0) | 20 (0) | Sub 47: L>F; Sub 49: K>T; Sub 50: N>K |
| BVQ_09750 | | | 7 (0) | 7 (0) | 7 (0) | 7 (0) | Sub 2141: A>S; Sub 2180: C>G |
| BVQ_09755 | | | 60 (0) | 60 (0) | 60 (0) | 60 (0) | Sub 546: E>D; Sub 691: A>S; Sub 698: A>G; Sub 717: A>E; Sub 753: V>M; Sub 797: L>F; Sub 836: N>D; Sub 901: T>N; Sub 905: E>D; Sub 930: A>V; Sub 940: M>V; Sub 1277: R>G |
| BVQ_09765 | | | 39 (0) | 38 (0) | 38 (0) | 38 (0) | Sub 561: S>A; Sub 568: P>S; Sub 584: S>G; Sub 593: E>D; Sub 597: A>S; Sub 961: R>Q; Sub 1139: |

| | | | | | | | |
|---|---|---|---|---|---|---|---|
| | | | | | | | P>A; Sub 1147: C>G; Sub 1272: R>G |
| BVQ_09930 | | | 12 (0) | 12 (0) | 12 (0) | 12 (0) | Sub 372: I>V; Sub 375: A>V |
| BVQ_10235 | | | 2 (0) | | | | Sub 72: L>I |
| BVQ_10270 | | | | 1 (0) | | | Sub 193: G>D |
| BVQ_11585 | yunB | Sporulation protein YunB | 16 (0) | 16 (0) | 16 (0) | 16 (0) | Sub 15: M>I; Sub 105: V>A; Sub 117: G>D |
| BVQ_11600 | | | 3 (0) | 3 (0) | 3 (0) | 3 (0) | No differences |
| BVQ_12510 | | | 16 (0) | 16 (0) | 16 (0) | 16 (0) | Sub 3134: V>I; Sub 3155: F>V; Sub 3161: S>F; Sub 3169: I>V; Sub 3180: E>D; Sub 3199: S>P; Sub 3204: G>D |
| BVQ_12525 | | | 5 (0) | 5 (0) | 5 (0) | 5 (0) | Sub 3005: D>G; Sub 3037: C>G; Sub 3110: D>G; Sub 3231: A>T |
| BVQ_13970 | | | 5 (0) | 5 (0) | 5 (0) | 5 (0) | No differences |
| BVQ_14470 | | | 7 (0) | 7 (0) | 7 (0) | 7 (0) | Sub 3: R>G; Sub 9: S>C ; Sub 14: G>R; Sub 28: R>G; Sub 53: F>S; Sub 53: F>S; Sub 55: G>R; Sub 62: T>A; |
| BVQ_16310 | | | 1 (0) | 1 (0) | 1 (0) | 1 (0) | Sub 170: R>H |
| BVQ_17710 | | | 1 (0) | | | | Sub 526: N>T |
| BVQ_18420 | | | 106 (2) | 106 (2) | 106 (2) | 106 (2) | Sub 850: S>V; Sub 853: A>S; Sub 924: Q>K; Sub 938: G>E; Sub 939: E>G; Sub 940: T>A; Sub 941: L>Y; Sub 1030: I>L; Sub 1041: G>A; Sub 1049: N>D; Sub 1087: K>R; Sub 1091: V>A; Sub 1094: E>Q; Sub 1100: N>Q; Sub 1212: D>N; Sub 1214: D>E; Sub 1225: D>E; Sub 1230: H>P; Sub 1245: H>Y; Sub 1262: Q>E; Sub 1284: D>E; Sub 1291: H>Y; Sub 1307: N>D; Del 1312-1314: HTE; Sub 1318: H>N; Sub 1319: H>N; Sub 1376: A>T; Sub 1377: S>T; Sub 1379: G>E; Sub 1395: S>A; Sub 2274: Y>H; Sub 2291: E>Q; Sub 2313: E>D; Sub 2320: Y>H; Sub 2336: D>N; Ins 2341-2343: HTE; Sub 2347: N>H; Sub 2348: N>H; Sub 2833: L>P |
| BVQ_19105 | | | | | 1 (0) | | No differences |

**[1]Sub = substitution, Ins = insertion, Del = deletion**

**Table S11: Specialised metabolites specifically targeted by metabolic profiling using HPLC-HRMS, based on genome mining and antiSMASH analysis (see Figure 6 in the manuscript) that were not produced by**

**any of the three *B. velezensis* strains in the given cultivation conditions on LB agar (i.e., < LOD (Limit of detection)).**

| Compound name | Chemical formula | Exact mass | Chemical structure |
|---|---|---|---|
| **Butirosin A** | $C_{20}H_{39}N_5O_{12}$ | 541.25952 | 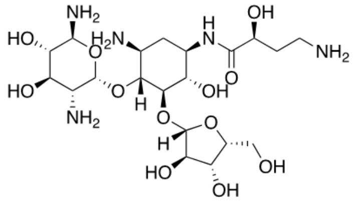 |
| **Butirosin B** | $C_{20}H_{39}N_5O_{12}$ | 541.25952 | 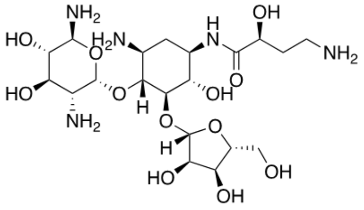 |
| **Citrulline** | $C_6H_{13}N_3O_3$ | 175.09569 | 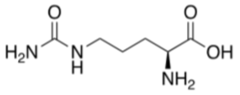 |
| **Macrolactin A** | $C_{24}H_{34}O_5$ | 402.24062 | 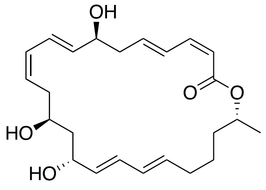 |
| **7-*O*-succinyl-macrolactin A** | $C_{28}H_{38}O_8$ | 502.25667 | 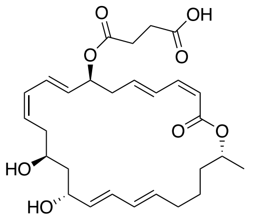 |
| **7-O-malonyl-macrolactin A** | $C_{27}H_{36}O_8$ | 488.24102 | 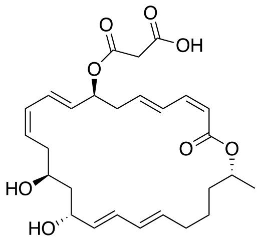 |
| **Macrolactin B** | $C_{30}H_{44}O_{10}$ | 564.29345 | 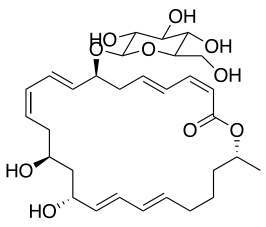 |

| | | | |
|---|---|---|---|
| **Macrolactin C** | $C_{30}H_{44}O_{10}$ | 564.29345 | 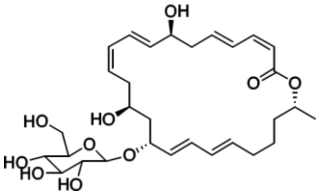 |
| **Macrolactin D** | $C_{34}H_{48}O_{13}$ | 664.30949 | 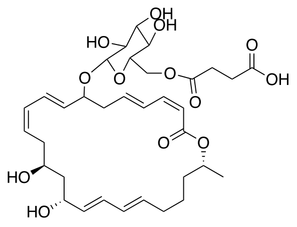 |
| **Macrolactin E** | $C_{24}H_{32}O_5$ | 400.22497 | 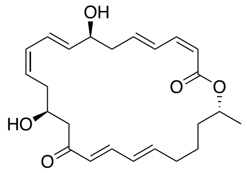 |
| **7-O-succinyl-macrolactin E** | $C_{28}H_{36}O_8$ | 500.24102 | 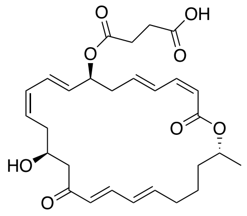 |
| **Macrolactin F** | $C_{24}H_{34}O_5$ | 402.24062 | 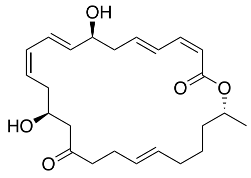 |
| **7-O-succinyl-macrolactin F** | $C_{28}H_{38}O_8$ | 502.25667 | 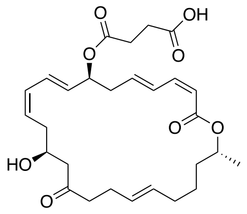 |
| **15-epi-dihydromacrolactin F** | $C_{24}H_{38}O_5$ | 406.27192 | 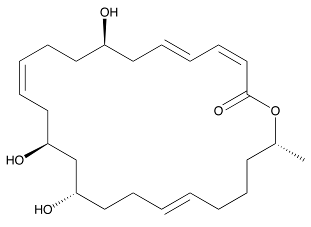 |
| **Macrolactin G** | $C_{24}H_{34}O_5$ | 402.24062 | 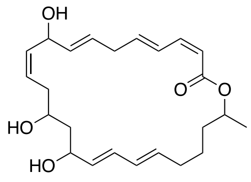 |

| | | | |
|---|---|---|---|
| **Macrolactin H** | $C_{22}H_{32}O_5$ | 376.22497 | 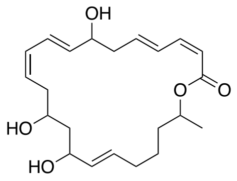 |
| **Macrolactin I** | $C_{24}H_{34}O_5$ | 402.24062 | 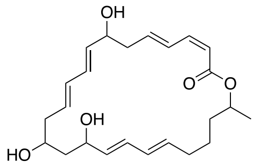 |
| **Macrolactin J** | $C_{24}H_{34}O_5$ | 402.24062 | 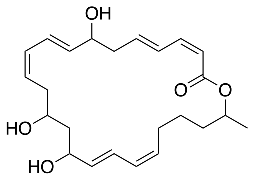 |
| **Macrolactin K** | $C_{24}H_{34}O_5$ | 402.24062 | 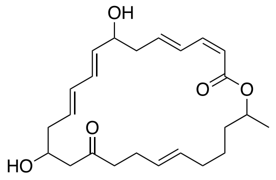 |
| **Macrolactin L** | $C_{24}H_{34}O_5$ | 402.24062 | 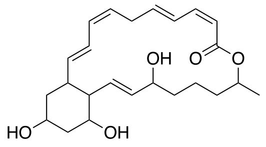 |
| **Macrolactin M** | $C_{25}H_{36}O_5$ | 416.25627 | 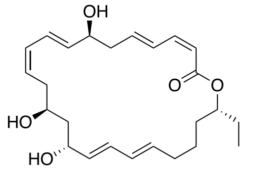 |
| **Macrolactin N** | $C_{24}H_{34}O_4$ | 386.24571 | 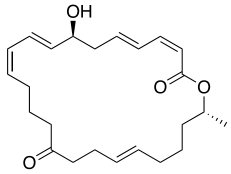 |
| **Macrolactin O** | $C_{30}H_{44}O_{10}$ | 564.29345 | 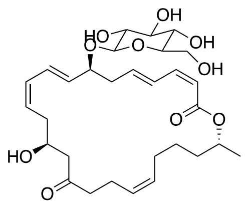 |
| **Macrolactin P** | $C_{31}H_{46}O_{10}$ | 578.30910 | 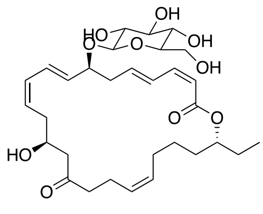 |

| | | | |
|---|---|---|---|
| **Macrolactin Q** | $C_{30}H_{44}O_{10}$ | 564.29345 | 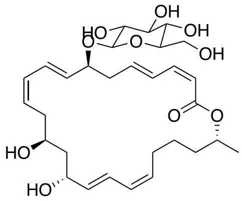 |
| **Macrolactin R** | $C_{30}H_{44}O_{10}$ | 564.29345 | 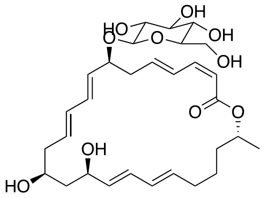 |
| **Macrolactin S** | $C_{24}H_{34}O_6$ | 418.23554 | 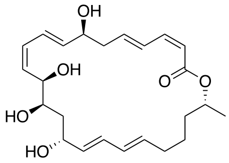 |
| **Macrolactin T** | $C_{24}H_{38}O_6$ | 422.26684 | 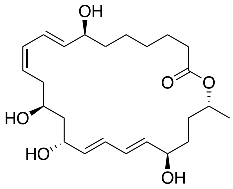 |
| **Macrolactin U** | $C_{31}H_{44}O_4$ | 480.32396 | 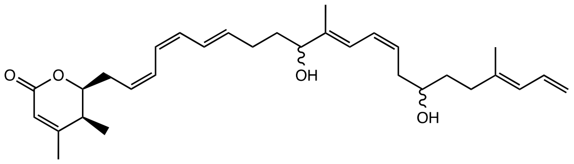 |
| **Macrolactin X** | $C_{24}H_{34}O_6$ | 418.23554 | 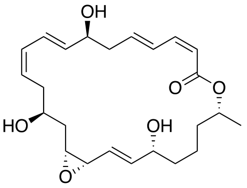 |
| **Macrolactin Y** | $C_{25}H_{38}O_7$ | 450.26175 | 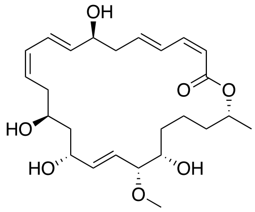 |
| **Macrolactin Z** | $C_{29}H_{40}O_8$ | 516.27232 | 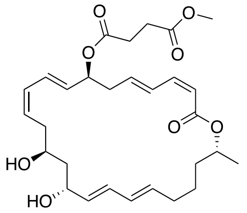 |

| | | | |
|---|---|---|---|
| **7-o-methyl-5-hydroxy-3-heptonate-macrolactin** | $C_{31}H_{48}O_7$ | 532.340 | 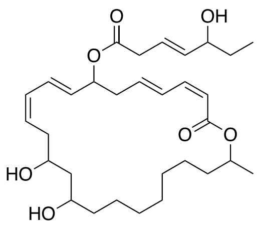 |

#### Supplementary Figures

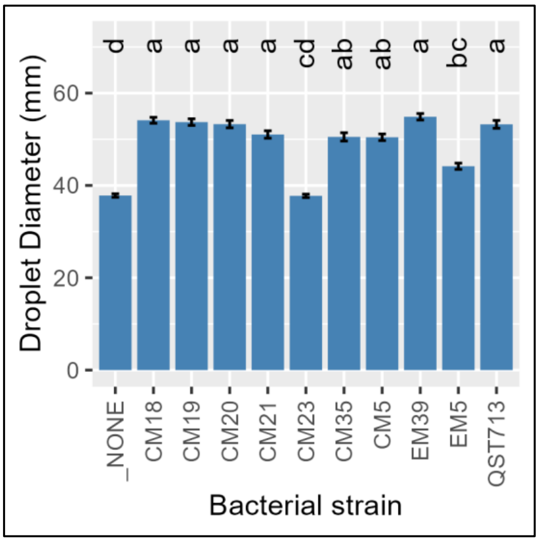


**Figure S1: Surfactant production in *B. velezensis* strains. Data shown is the mean droplet diameter of supernatant from overnight LB cultures. Droplet diameters were measured 2 minutes after pipetting onto microscope slides. Graph shows the mean and standard error of 5 experiments, each containing 5 droplets per treatment. Letters indicate groups of statistically different conditions (one-way ANOVA and Tukey's HSD test α = 0.05).**
